# RasGEFX triggers spontaneous Ras excitation with RasGEFB/M/U for random cell migration

**DOI:** 10.1101/2023.08.13.553116

**Authors:** Koji Iwamoto, Satomi Matsuoka, Masahiro Ueda

**Affiliations:** Laboratory of Single Molecule Biology, Graduate School of Science and Graduate School of Frontier Biosciences, Osaka University, 1-3 Yamadaoka, Suita, Osaka 565-0871 Japan; Laboratory for Cell Signaling Dynamics, Center for Biosystems Dynamics Research (BDR), RIKEN, 1-3 Yamadaoka, Suita, Osaka 565-0871; PRESTO, JST, 1-3 Yamadaoka, Suita, Osaka 565-0871 Japan

**Keywords:** Ras, RasGEF, excitable system, chemotaxis, *Dictyostelium*, hierarchical clustering, spontaneous activity, positive feedback

## Abstract

Excitable systems of eukaryotic chemotaxis can generate asymmetric signals of Ras-GTP-enriched domains spontaneously to drive random cell migration without guidance cues. The molecules responsible for the spontaneous signal generation remain elusive. Here, we characterized RasGEFs encoded in *Dictyostelium discoideum* by the live-cell imaging of the Ras dynamics and hierarchical clustering, finding that RasGEFX triggers symmetry breaking to generate a Ras-GTP-enriched domain and is essential for random migration in combination with RasGEFB/M/U but dispensable for chemotaxis. Among RasGEFs, RasGEFX and RasGEFB co-localize with Ras-GTP on oscillatory waves propagating along the membrane and regulate the temporal periods and spatial sizes, respectively, of the waves and thus the cytoskeletal dynamics differently. RasGEFU localized uniformly on the membrane and regulated cell adhesion. These findings provide mechanistic insights into the internal signal generation that operates independently of external chemotaxis signaling and suggest a coordinated control of the cytoskeletal dynamics by multiple RasGEFs for spontaneous motility.

## INTRODUCTION

Spontaneous activity is one of the hallmarks of living organisms. Spontaneous cell migration is a typical example of spontaneous activity and describes how various cells exhibit motile function in a random manner without environmental guidance cues, which is a basis for directed cell migration.^1–6^ Some specific molecular processes are likely to generate signals internally to drive cell motility. Recent studies have demonstrated that excitable systems of chemotaxis generate an asymmetric signal to regulate cell motility spontaneously by breaking symmetry in the intracellular distribution of various signaling molecules, including Ras, PI3K, PI(3,4,5)P3, Akt/PKB and Scar/WAVE at the “front” region of motile cells and PTEN, PI(4,5)P2 and CynA at the “back” region.^7–11^ Such excitable systems are conserved evolutionarily in the chemotactic signaling networks of various cell types from the lower eukaryote *Dictyostelium discoideum* to human leukocytes.^11–16^ However, our understanding of how spontaneous dynamics emerge from excitable systems is limited because the molecules responsible for triggering the symmetry breaking have not been identified.

Ras is a small GTPase and key molecule for cellular anterior-posterior polarity and motility, including chemotaxis, in various motile cells.^11,17^ In *Dictyostelium* cells, the active form of Ras, Ras-GTP, generates Ras-GTP/PI3K/PI(3,4,5)P3-enriched signaling domains to regulate cytoskeletal dynamics such as pseudopod formation.^4,8,18,19^ The signaling molecules-enriched domain exhibits stereotypical characteristics of an excitable system, including spontaneous excitation, stimulation-induced excitation and traveling wave generation.^11,13,14,20–24^ For example, the Ras-GTP-enriched domain is generated transiently on the plasma membrane in an all-or-none manner through spontaneous excitatory dynamics without the extracellular chemoattractant cAMP. At the same time, under enhanced excitability, the domain can exhibit traveling waves propagating continuously over the membrane as oscillatory dynamics.^24,25^ The transition between excitatory and oscillatory dynamics have been theoretically predicted as a characteristic of excitable systems and can be manipulated experimentally by treating *Dictyostelium* cells with caffeine.^24,26–28^ Further evidence for excitability includes chemoattractant-induced all-or-none excitation, in which the Ras-GTP-enriched domain is induced by a uniform stimulation of cAMP or biased spatially under cAMP gradients toward the higher concentration side.^23,29,30^ None of the four major signaling pathways that mediate chemoattractant signals downstream of Ras: the PI3K/PI(3,4,5)P3, TorC2, PLA2 and soluble guanylate cyclase (sGC) pathways, are required for the spontaneous generation of the Ras-GTP-enriched domain when cAMP is absent.^24,29–32^ That is, the Ras-GTP-enriched domain is generated without upstream signaling from chemoattractant receptors and without the activities of downstream pathways. Thus, identifying the molecules responsible for the Ras excitability is a key to understand the spontaneous symmetry breaking for cell motility.

Ras activity is regulated by guanine nucleotide exchange factors (RasGEFs) and GTPase-activating proteins (RasGAPs). So far, 25 genes for RasGEFs and 14 genes for RasGAPs have been found in the *Dictyostelium discoideum* genome,^33,34^ with several RasGEFs (A/F/H/Q/R/U) considered potential regulators of Ras.^35–40^ Upon cAMP binding to the receptor cAR1, downstream signaling via the heterotrimeric G protein Gα_2_βγ leads to the transient activation of Ras, in which RasGEFA and RasGEFH activate RasC, and RasGEFR and RasGEFF activate RasG.^37,39,40^ Additionally, RasGEFQ and RasGEFU activate RasB and Rap1 upon cAMP stimulation, respectively.^36,38^ However, the contributions of these and other RasGEFs to the spontaneous dynamics of excitable systems are unknown, because a systematic and comprehensive analysis of RasGEFs on Ras excitability without chemoattractants has never been performed.

In this study, we developed a screening method to identify the RasGEFs responsible for Ras excitability in *Dictyostelium* in the absence of cAMP by live-cell imaging and hierarchical clustering analysis. We found that specific RasGEFs break symmetry in the excitable system to generate internal signals spontaneously for random cell motility and that these RasGEFs can operate independently of external chemotaxis signaling. RasGEFX is primarily required for the spontaneous symmetry breaking and thus for basal cell motility. However, RasGEFB/M/U regulate Ras excitability and are required in combination with RasGEFX for random cell motility. From these observations, we propose that multiple RasGEFs constitute a spontaneous signal generator in a concerted manner to drive random cell motility.

## RESULTS

### Comprehensive characterization of RasGEFs in Ras excitability

In order to identify the RasGEFs responsible for spontaneous excitation of the Ras excitable system, we characterized the spatiotemporal dynamics of the Ras-GTP-enriched domain when RasGEFs were overexpressed. We prepared a series of cell strains overexpressing (OE) one of 25 RasGEFs tagged to GFP and co-expressed them with RBD_Raf1_-RFP, a fluorescent reporter of Ras-GTP, for live-cell imaging of the Ras-GTP-enriched domain^18,41^ (Figures 1A and 1B). When introduced in wild-type Ax2 cells, 22 RasGEF OE strains were stable for the expression of RBD_Raf1_-RFP. The other 3 transformants (RasGEFI/J/Q) were unstable and thus excluded from the subsequent analysis. The remaining cells were treated with 10 μM latrunculin A, an inhibitor of actin polymerization, to exclude the effects of the cell shape on the Ras-GTP distribution. The cells were also treated with 4 mM caffeine to induce the oscillatory dynamics, in which the traveling waves of the Ras-GTP-enriched domain were observed, allowing us to compare the effects of each OE strain on the spatiotemporal dynamics of Ras-GTP (Video S1). Under these conditions, we observed four patterns in the membrane localization of RBD_Raf1_-RFP in wildtype cells (WT cells): no generation of the Ras-GTP-enriched domain (no domain), transient generation of the Ras-GTP-enriched domain (transient), traveling waves (wave), and uniform localization along the whole membrane (uniform) (Figure 1C; Video S1). These patterns were observed for 20 min within individual cells and distinguished by kymographs of the RBD_Raf1_-RFP intensity measured along the cell periphery and by a hierarchical clustering analysis^42^ (Figures S1A-S1F). The fractions of the patterns in WT cells were 25.5 ± 5.4% (no domain), 13.8 ± 6.3% (transient), 57.2 ± 2.5% (wave) and 3.4 ± 1.7% (uniform) (Figure 1D; Table S1). The domain size and the oscillation period of the traveling waves in WT cells were 139.1 ± 61.0° (*n*=163 cells) and 244.0 ± 81.8 sec (*n*=124 cells), respectively (Figure S1G and S1H; Table S1). As shown in the subsequent analysis, these characteristics of the spatiotemporal dynamics of the Ras-GTP-enriched domain differed depending on the type of RasGEF overexpressed, allowing us to screen for the RasGEFs responsible for spontaneous Ras excitation.

**Figure 1.**
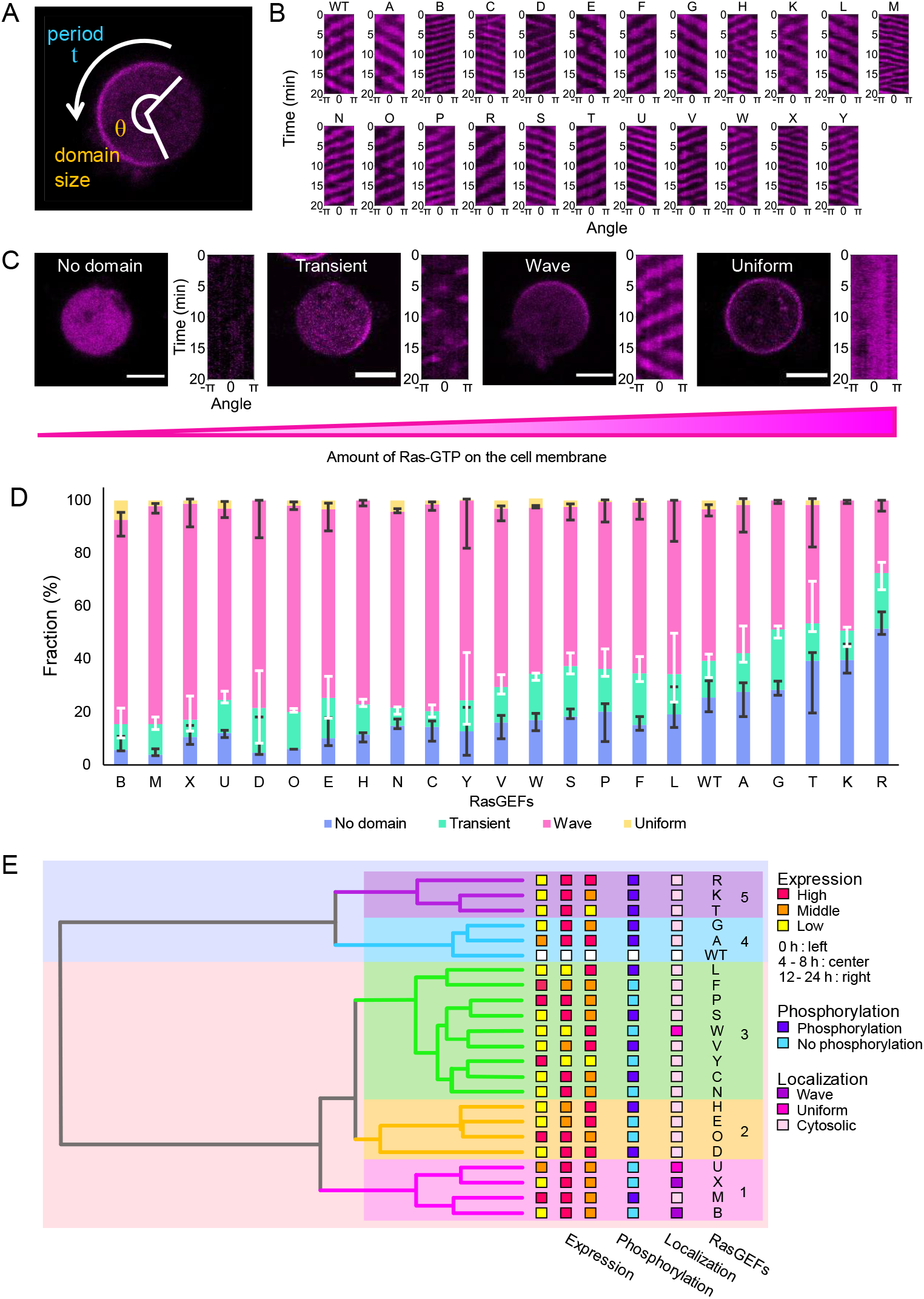
Screening for RasGEFs essential for the spontaneous excitability of Ras. (A) An image of a *Dictyostelium discoideum* cell expressing RBD_Raf1_-RFP. The spontaneous dynamics of Ras-GTP was analyzed by measuring the fluorescence intensity along the cell membrane over time to generate a kymograph. (B) Representative kymographs showing traveling waves of Ras-GTP in RasGEF OE strains. (C) Four patterns in the membrane localization of RBD_Raf1_-RFP. The excitability increases from left to right as the amount of Ras-GTP on the cell membrane increases. Scale bars, 5 μm. (D) Fraction of cells showing each pattern: no domain (*blue*), transient domain (*green*), traveling wave (*magenta*), and uniform (*yellow*). *n* > 110 cells from 2∼3 independent experiments for each strain. (E) Hierarchical clustering of RasGEFs based on the spontaneous dynamics of Ras-GTP in OE cells. Clusters 1 (*bottom*) through 5 (*top*) are arranged in order of the extent of Ras excitability, with cluster 1 the highest. Colored squares indicate the features of RasGEFs in wild-type cells: high (*red*), middle (*orange*) or low (*yellow*) expression levels at the indicated time windows during development; phosphorylated (*blue*) or not phosphorylated (*pale blue*) upon cAMP stimulation; localized on the cell membrane with traveling waves (*violet*), uniformly on the cell membrane (*pink*), or throughout the cytoplasm (*pale pink*).

We found that the overexpression of a RasGEF acts either positively or negatively on Ras excitability. When compared with WT cells, the fraction of traveling waves was increased in the majority of RasGEF OE strains (B/M/X/U/D/O/E/H/N/C/Y/V/W/S/P/F/L) but decreased in the others (A/G/T/K/R) (Figure 1D). The fraction of cell sub-populations with no domain generation varied depending on the overexpressed RasGEF. The fraction was negatively correlated with traveling wave fraction, showing that wave generation is an indicator of the degree of excitability under caffeine-treated conditions. The mean domain size was larger in only the RasGEFB OE strain but smaller in many more (A/D/G/H/K/L/O/P/R/Y) compared with WT cells (Figure S1G; Table S1). The mean period was shorter in the RasGEF OE strains (B/C/D/H/M/N/O/P/S/T/U/V/W/X/Y), and no strains showed a longer period compared with WT cells (Figure S1H; Table S1).

We classified the 22 RasGEFs by hierarchical clustering into clusters for which overexpression altered the Ras dynamics in a similar manner (Figures 1D, 1E and S2). The domain sizes and oscillatory periods of the Ras-GTP-enriched domain were determined from individual RasGEF OE cells (Figures S1G and S1H), and corresponding heatmaps were obtained for each cell population (Figure S2). Based on an analysis of the heatmap, RasGEFs were grouped into five clusters and aligned so that the sum of both the fractions of the traveling wave and uniform domain subpopulations increased from top to bottom (Figures 1D and 1E). Clusters 1 and 5 showed the largest positive and negative effects on Ras excitability, respectively (Figure 1E). RasGEFB/M/X/U in cluster 1 enhanced the excitability relatively strongly, while RasGEFT/K/R in cluster 5 suppressed the excitability. In cluster 1, endogenous RasGEFs showed a peak in their gene expressions at around 4 to 8 hours after starvation.^43^ Furthermore, cAMP did not phosphorylate them,^44^ and they localized to the cell membrane even without cAMP or functional F-actin, except for RasGEFM, as shown below. In cluster 5, endogenous RasGEFs also showed a peak in gene expressions at around 4 to 8 hours after starvation. However, they were phosphorylated by chemoattractant stimulation, suggesting their involvement in chemotactic signaling. RasGEFT/K/R hardly localized to the cell membrane without cAMP or functional F-actin, as shown below. RasGEFs in cluster 4 shared similar characteristics to those in cluster 5, whereas the RasGEFs in clusters 2 and 3 shared few common characteristics. Considering that *Dictyostelium* cells can exhibit random migration without cAMP and directional migration with cAMP extensively after 4 hours of starvation,^6^ we concluded that RasGEFs in cluster 1 likely promote spontaneous excitability, while those in clusters 4 and 5 likely suppress spontaneous excitability but may be involved in the response to cAMP through chemical modifications such as phosphorylation.

### RasGEFX is primarily required for Ras excitability

To elucidate the contributions of the four RasGEFs in cluster 1 on the excitable dynamics of Ras, we prepared 4 single knock-out (KO) strains and 6 double knock-out (DKO) stains by CRISPR/Cas9 or homologous recombination, in which mScarlet-RBD was expressed to observe the Ras-GTP-enriched domain^45^ (Figure 2A; Video S2). In conditions where excitability was enhanced with caffeine, no *gefX*-cells showed traveling waves (n=286 cells), while 66.1 ± 5.2% of WT cells (n=200 cells) did (Figure 2B; Table S1). The fraction of KO cells showing traveling waves was 85.2 ± 6.3% (n=191 cells), 77.6 ± 1.9% (n=210 cells) and 73.4 ± 4.3% (n=217 cells) by disrupting *gefB, gefM* and *gefU*, respectively, but they were reduced to almost zero with the DKO of *gefX* (Figure 2B). The DKOs with other combinations of RasGEFB/M/U exhibited significant wave generation, with *gefB-/M-* and *gefB-/U-* cells especially exhibiting enhanced excitability (Figure 2B). These results indicate that RasGEFX is required for the generation of traveling waves in the presence of caffeine and that RasGEFB/M/U are suppressive of Ras excitability even though their overexpression enhances it (Figure 1). These apparently inconsistent observations can be explained by RasGEFB/M/U acting competitively against RasGEFX in the activation of Ras and having weaker GEF activity than RasGEFX. Therefore, it is likely that RasGEFX fully enhances excitability in cells in which competitive *gefB, gefM*, and *gefU* are knocked out.

**Figure 2.**
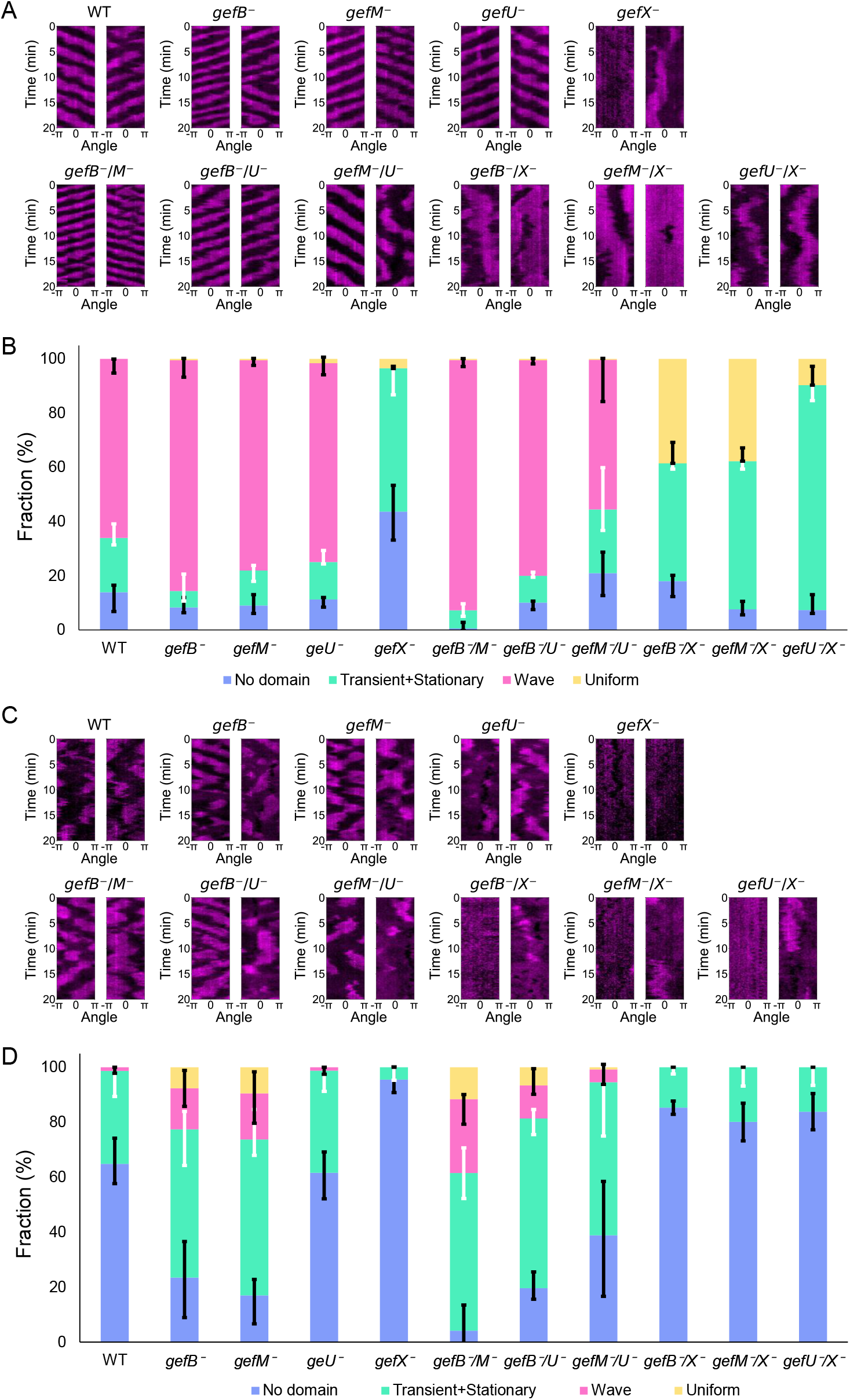
RasGEFX is primarily required for Ras excitability. (A, B) Ras-GTP dynamics in RasGEF KO strains in the presence of caffeine. (A) Representative kymographs of mScarlet-RBD_Raf1_. (B) Fraction of cells showing each pattern: no domain (*blue*), transient domain and stationary domain (non-oscillatory domains; *green*), traveling wave (*magenta*), uniform (*yellow*). The means and SDs of 2 or 3 independent experiments are shown (WT, *n* = 200 cells; *gefB-, n* = 191 cells; *gefM-, n* = 210 cells; *gefU-, n* = 217 cells; *gefX-, n* = 286 cells; *gefB-/M-, n* = 213 cells; *gefB-/U-, n* = 223 cells; *gefM-/U-, n* = 314 cells; *gefB-/X-, n* = 202 cells; *gefM-/X-, n* = 225 cells; *gefU-/X-, n* = 271 cells). (C, D) Ras-GTP dynamics in RasGEF KO strains in the absence of caffeine. (C) Representative kymographs of mScarlet-RBD_Raf1_. (D) Fraction of cells showing each pattern: no domain (*blue*), non-oscillatory domains (*green*), traveling wave (*magenta*), uniform (*yellow*). The means and SDs of 3 to 6 independent experiments are shown (WT, *n* = 87 cells; *gefB-, n* = 105 cells; *gefM-, n* = 119 cells; *gefU-, n* = 89 cells; *gefX-, n* = 121 cells; *gefB-/M-, n* = 71 cells; *gefB-/U-, n* = 85 cells; *gefM-/U-, n* = 107 cells; *gefB-/X-, n* = 87 cells; *gefM-/X-, n* = 123 cells; *gefU-/X-, n* = 123 cells).

We then observed the spontaneous Ras-GTP dynamics in the same KO and OE strains under physiological conditions where excitability is not enhanced by caffeine. WT cells exhibited a transient generation of the Ras-GTP-enriched domain (33.9 ± 9.4%, n=87 cells) but little or no generation of a traveling wave (Figures 2C and 2D; Table S1; Video S2). As expected, RasGEFB/M/U/X OE cells enhanced Ras excitability even in the absence of caffeine, showing the excitatory roles of cluster-1 RasGEFs when highly expressed (Figures S3A and S3B). Cells missing RasGEFX (*gefX-, gefB-/X-, gefU-/X-* and *gefM-/X-*) showed impaired domain generation, consistent with the caffeine observations (Figures 2C and 2D; Table S1; Video S2). On the other hand, the disruption of RasGEFB/M/U (i.e. *gefB-, gefM-, gefB-/M-, gefB-/U-* and *gefM-/U-*), in which RasGEFX is expected to be intact, tended to increase the Ras excitability, in which traveling waves were often observed even in the absence of caffeine. These results indicate that RasGEFX is required for the spontaneous excitation to generate the Ras-GTP-enriched domain under physiological conditions, while RasGEFB/M/U are competitive with RasGEFX, probably due to their weaker RasGEF activity, but can enhance the spontaneous excitability with their exogenous expression. Thus, RasGEFX is primarily responsible for spontaneous symmetry breaking in the Ras excitable system to generate an asymmetric Ras-GTP-enriched domain, while RasGEFB/M/U modulate the spatiotemporal dynamics.

### RasGEFX is required for random cell motility in combination with RasGEFB/M/U

To evaluate the involvement of cluster-1 RasGEFB/M/U/X in spontaneous cell motility, we examined the motility of KO and OE cells by analyzing the migration trajectories in the absence of both caffeine and a chemoattractant. No RasGEFB/M/U/X KO cells showed serious defects in cell motility, although the sub-population of *gefX-* cells exhibited almost no migration (Figures 3A and 3B; Video S3). The mean migration speeds were 7.0 ± 3.5 μm/min for WT cells (n=77 cells), 4.4 ± 3.5 μm/min for *gefX-* (n=65 cells), 5.5 ± 2.9 μm/min for *gefM-* (n=77 cells), 10.6 ± 4.8 μm/min for *gefB-* (n=67 cells) and 10.0 ± 3.4 μm/min for *gefU-* (n=70 cells (Figure 3B; Table S1). RasGEFX DKO cells, including *gefB-/X-, gefU-/X-* and *gefM-/X-*, showed severely impaired spontaneous cell migration, whereas other DKO cells, including *gefB-/M-, gefB-/U-* and *gefM-/U-*, exhibited spontaneous migration, albeit with slight defects (Figures 3A and 3B). The mean square displacement (MSD) of the migration trajectories confirmed that the spontaneous cell motility depended on RasGEFX in combination with RasGEFB/M/U (Figure 3C). The migration speed decreased in all OE strains compared to WT cells (Figures 3D-3F; Table S1; Video S3) due to the abnormal formation of pseudopods over the whole cell surface by over-activation of the excitable system. In addition, the overexpression of RasGEFU enhanced cell adhesion to substrates, consistent with previous reports of cGMP-binding protein D (GbpD), which is the same as RasGEFU^36,46^ (Figure S3C). The hierarchical clustering analysis based on the dynamics of Ras-GTP with caffeine showed that the genetic disruption of RasGEFX (i.e. gefX-, gefB-/X-, gefU-/X- and gefM-/X-) caused a significant decrease in the traveling wave generation; thus, we separated these four strains from the other KO strains (Figures 3G and S3D). Even in the absence of caffeine, the cluster in which RasGEFX gene was disrupted showed impaired domain generation. Furthermore, the DKOs in this cluster exhibited a significant decrease in the migration speed. Conversely, the strains in the other cluster with intact RasGEFX exhibited relatively higher motility. These results indicate that RasGEFX plays a key role in the spontaneous signal generation for basal cell motility under no extracellular signals. They also indicate that the other RasGEFs (B/M/U) in cluster 1 are required with RasGEFX to achieve normal cell migration.

**Figure 3.**
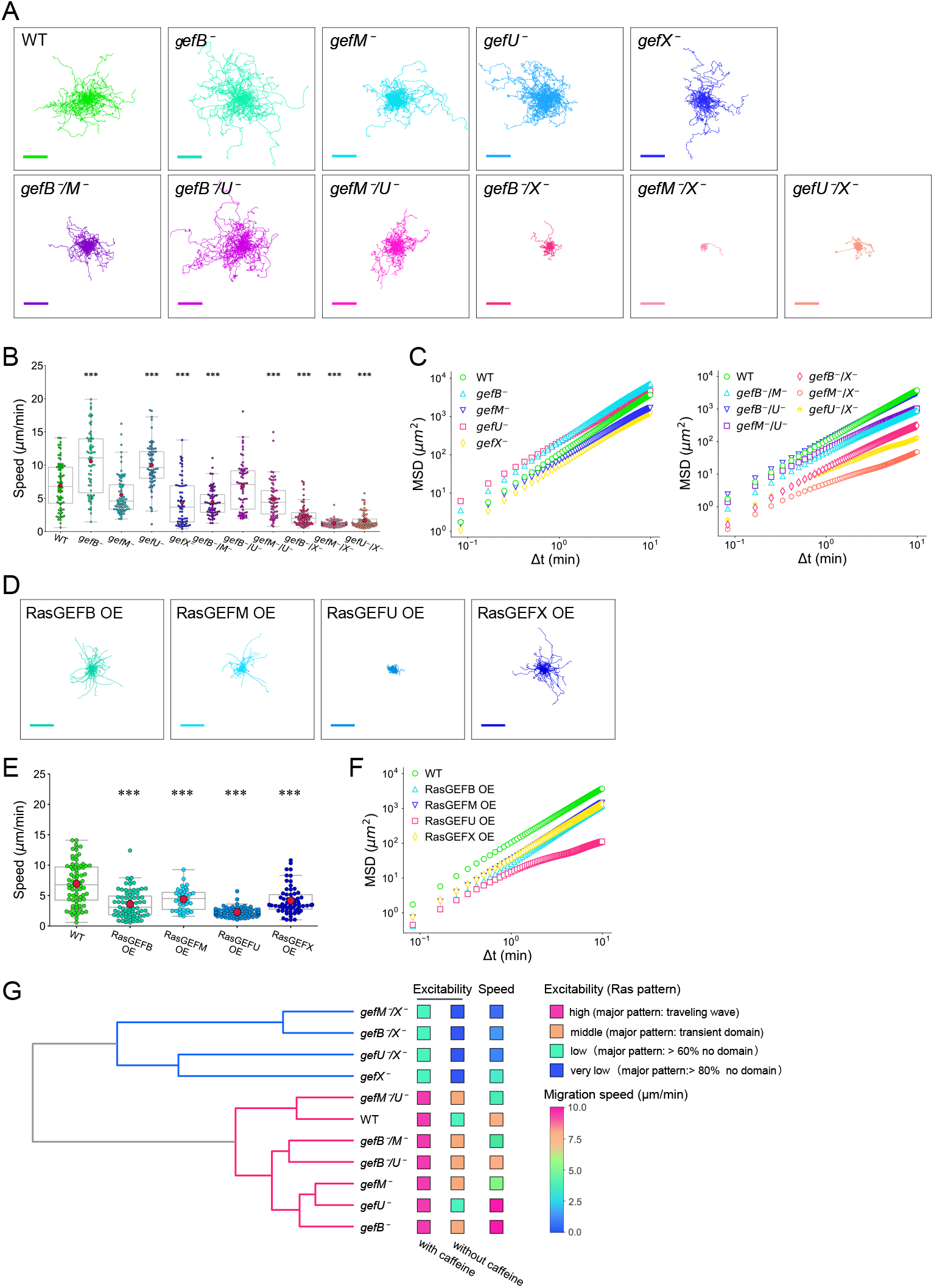
RasGEFX, RasGEFB, RasGEFU and RasGEFM are important for efficient spontaneous motility. (A-C) Statistical analysis of spontaneous motility by RasGEF KO strains. (A) Trajectories of spontaneously migrating KO cells. WT, *n* = 77 cells; *gefB-, n* = 67 cells; *gefM-, n* = 77 cells; *gefU-, n* = 70 cells; *gefX-, n* = 65 cells; *gefB-/M-, n* = 66 cells; *gefB-/U-, n* = 66 cells; *gefM-/U-, n* = 62 cells; *gefB-/X-, n* = 70 cells; *gefM-/X-, n* = 65 cells; *gefU-/X-, n* = 66 cells. Scale bars, 100 μm. (B) Speed of spontaneous migration. (C) MSD calculated for each KO strain. (D-F) Statistical analysis of spontaneous motility by RasGEF OE strains. (D) Trajectories of spontaneously migrating OE cells. RasGEFB OE, *n* = 75 cells; RasGEFM OE, *n* = 39 cells; RasGEFU OE, *n* = 66 cells; RasGEFX OE, *n* = 65 cells. Scale bars, 100 μm. (E) Speed of the spontaneous migration. (F) MSD calculated for each OE strain. (B and E) Closed circles in magenta show the mean values. (G) Hierarchical clustering of RasGEF KO strains based on the spontaneous dynamics of Ras-GTP with caffeine. Clusters 1 (*bottom*) and 2 (*top*) are arranged in order of the extent of Ras excitability, with cluster 1 high. Colored squares indicate the features of RasGEF KO strains: high (*pink*), middle (*orange*), low (*cyan*) or very low (*blue*) fraction of domain patterns; migration speed is referred to the color bar. * *p* < 0.05, ** *p* < 0.01, *** *p* < 0.001 as calculated using Dunnett’s test.

### RasGEFX and RasGEFB co-localize in the Ras-GTP-enriched domain to generate traveling waves

To reveal the mechanisms by which cluster-1 RasGEFs underlie and regulate Ras excitability, we examined the subcellular localization of GFP-tagged RasGEFs. Among the 25 RasGEFs, only RasGEFB/U/X in cluster 1 and RasGEFW in cluster 3 exhibited obvious localization to the plasma membrane (Figure S4). RasGEFB/U/X have no transmembrane domain, suggesting dynamic shuttling between the membrane and the cytosol; in contrast, RasGEFW has a transmembrane domain.^34^ Among cluster-1 RasGEFs, RasGEFX and RasGEFB exhibited traveling waves co-localizing with Ras-GTP, and cross-correlation functions between each RasGEF and Ras-GTP revealed a tight coincidence of localization and oscillatory dynamics with almost no time lag (Figures 4A and 4B; Video S4), suggesting their direct involvement in traveling wave generation. Consistently, RasGEFB/X-GFP were observed in the cytosol but not on the membrane when the cells had no Ras-GTP-enriched domain. RasGEFU localized almost uniformly to the membrane and did not exhibit traveling waves, while RasGEFM was observed throughout the cytoplasm (Figures 4C and 4D; Video S4), suggesting uniform modulation of GEF activity on the membrane and in the cytosol by RasGEFU and RasGEFM, respectively.

**Figure 4.**
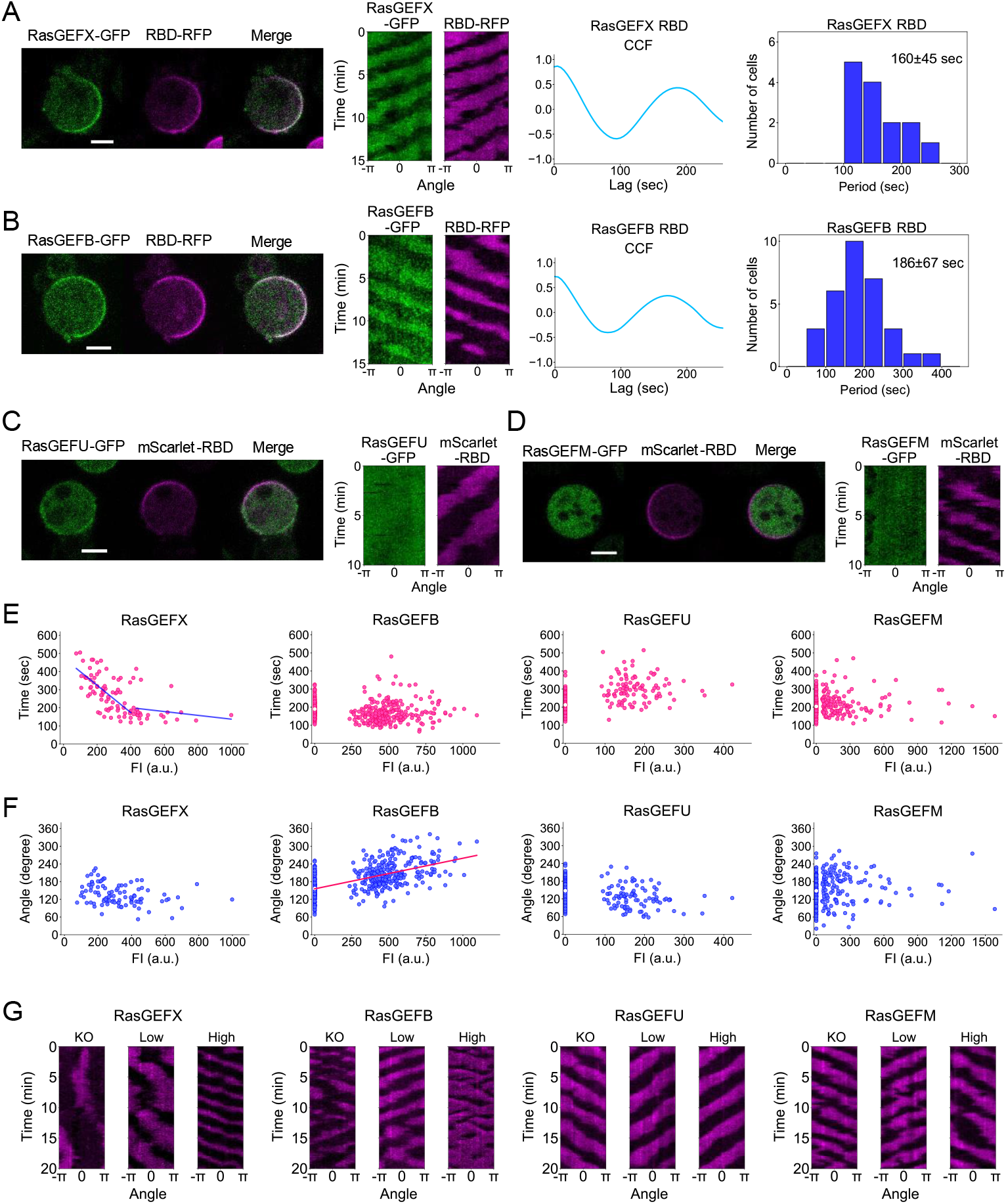
RasGEFX and RasGEFB co-localize with the traveling waves of Ras-GTP. (A-D) Co-localization analysis of RasGEFX (A), RasGEFB (B), RasGEFU (C) and RasGEFM (D) with Ras-GTP. (A and B) Representative images of RasGEFX-GFP (A) and RasGEFB-GFP (B) with RBD_Raf1_-RFP expressed in the respective RasGEF KO strains (*left*); representative kymographs (*middle left*); cross-correlation functions (CCFs) between RasGEF and RBD_Raf1_ (*middle right*); and the statistical distributions of the period estimated using the CCFs (*right*). RasGEFX, *n* = 15 cells; RasGEFB, *n* = 32 cells. (C and D) Representative images of RasGEFU-GFP (C) and RasGEFM-GFP (D) with mScarlet-RBD_Raf1_ expressed in the respective RasGEF KO strain (*left*) and kymographs (*right*). (E and F) Relationships between the period (E) and domain size (F) of Ras-GTP-enriched domain with the expression level of RasGEF, quantified in the RasGEF KO strains with (FI > 0) or without RasGEF-GFP expression (FI = 0). Closed circles represent the quantification in a single cell (*gefX-, n* = 97 cells; *gefB-, n* = 173 cells; *gefU-, n* = 101 cells; *gefM-, n* = 132 cells). (G) Representative kymographs of RBD_Raf1_-RFP or mScarlet-RBD_Raf1_ without (“KO”), with low (“Low”), or high expression (“High”) of RasGEF-GFP in the respective RasGEF KO cells. Scale bars, 5 μm.

To further examine how cluster-1 RasGEFs regulate the spatiotemporal dynamics of the Ras-GTP-enriched domain, the oscillation periods and the spatial sizes of the traveling waves were characterized by the expression level of RasGEFB/M/U/X. We prepared RasGEF KO strains expressing the corresponding GFP-tagged RasGEFs. Increasing the expression of RasGEFX-GFP shortened the oscillation period of the Ras excitable system but without any obvious correlation with the spatial size of the domain, indicating temporal regulation by RasGEFX (n=97 cells) (Figures 4E-4G). On the other hand, increasing the expression of RasGEFB-GFP resulted in larger domains with no obvious correlation with the oscillation period, indicating spatial regulation by RasGEFB (n=267 cells) (Figures 4E-4G). Neither the oscillation period nor the size was dependent on the expression level of RasGEFM (n=132 cells) or RasGEFU (n=101 cells) (Figures 4E-4G). These results indicate that RasGEFX and RasGEFB regulate the spatiotemporal dynamics of Ras excitability in different manners, such that the Ras-GTP-enriched domain can adopt diverse spatiotemporal dynamics as spontaneous excitability depending on the expression level ratio of the two RasGEFs. Because RasGEFX is responsible for the excitable firing and traveling wave generation of the Ras-GTP-enriched domain without and with caffeine, respectively, RasGEFX triggers spontaneous symmetry breaking as a temporal regulator in the excitable system to generate an asymmetric signal for basal cell motility. RasGEFB, on the other hand, modulates the spatial properties of the Ras-GTP-enriched domain in the excitable system. RasGEFM and RasGEFU hardly affect the spatiotemporal characteristics in Ras excitability in the absence of a functional actin cytoskeleton (i.e. presence of latrunculin A), but they can increase subpopulations of excited cells when they are overexpressed (Figure 1).

### RasGEFX and RasGEFB regulate actin cytoskeleton-dependent protrusion dynamics in a different manner

We further examined the subcellular localization of cluster-1 RasGEFs under the presence of a functional actin cytoskeleton without caffeine to reveal how the RasGEFs regulate actin cytoskeleton-dependent cellular processes such as pseudopod formation and micropinocytosis.^47,48^ As expected, RasGEFX and RasGEFB co-localized in the Ras-GTP-enriched domain at leading-edge pseudopods, while RasGEFU and RasGEFM showed uniform localization along the whole membrane and in the cytosol, respectively (Figure 5A; Video S5). That is, a functional actin cytoskeleton did not cause remarkable changes in the localization of cluster-1 RasGEFs. However, a characteristic morphological change was observed, especially in RasGEFX-GFP or RasGEFB-GFP-expressing cells, due to the cytoskeletal dynamics. RasGEFX-GFP-expressing cells often showed multiple pseudopods or macropinocytic cups in which RasGEFX-GFP were localized, and they frequently repeated the formation and retraction of such actin-dependent protrusions (Figure 5A). On the other hand, RasGEFB-GFP-expressing cells showed macropinocytosis less frequently than RasGEFX-GFP-expressing cells; instead, they often had multiple lamellipodia-like pseudopods extending continuously or one prominent larger pseudopod along the entire membrane (Figure 5A). RasGEFU-GFP-expressing cells showed increased adhesion to the substrates and thus less motility as their expression increased, and RasGEFM-GFP-expressing cells showed no changes compared to WT cells (Figure 5A).

**Figure 5.**
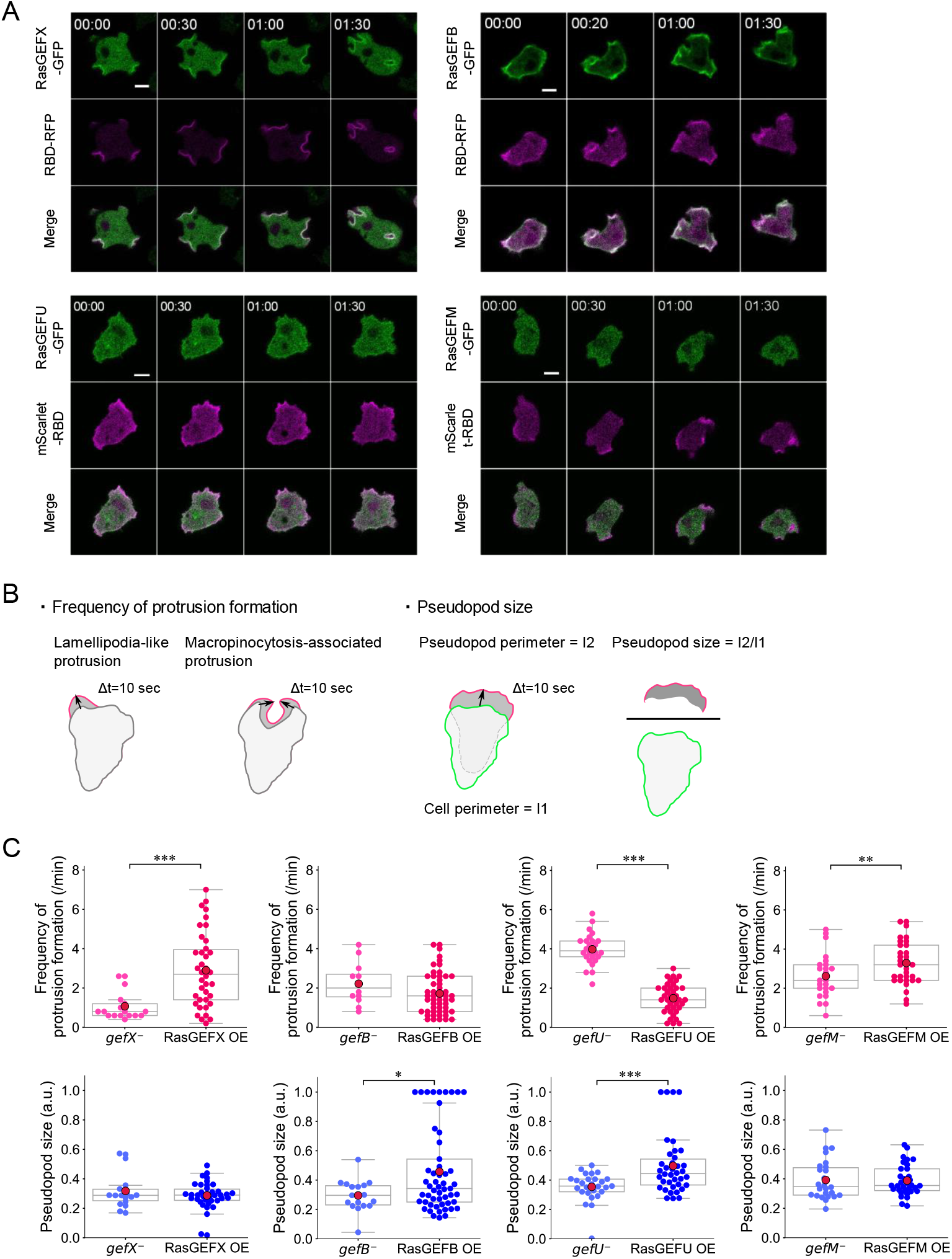
RasGEFX and RasGEFB are respectively responsible for the temporal and spatial dynamics of protrusions. (A) Representative images of RasGEF KO cells expressing RasGEF-GFP and RBD_Raf1_-RFP or mScarlet-RBD_Raf1_ with intact F-actin. Scale bars, 5 μm. Time, min:sec. (B) Schematics for the quantification of the frequency of the protrusion formation (*left*) and pseudopod size (*right*). (C) The frequency of the protrusion formation (*top*) and pseudopod size (*bottom*) in RasGEF KO and OE cells. * *p* < 0.1, ** *p* < 0.05, *** *p* < 0.01 as calculated using Welch’s *t*-test.

Because RasGEFX and RasGEFB regulate the oscillations temporally and the sizes spatially of the Ras-GTP-enriched domains without an actin cytoskeleton, respectively, we next analyzed the frequency and size of the actin-dependent protrusion formation under physiological conditions, where we did not distinguish between lamellipodia-like protrusions and macropinocytosis-associated protrusions (Figure 5B). *gefX-* cells were suppressed in the protrusion formation, and the formed area was small (Figure 5C; Video S5). RasGEFX overexpression in *gefX-* cells markedly promoted protrusion formation, but the area was not significantly altered, which is consistent with RasGEFX acting as a temporal regulator of Ras excitation. In contract, the other RasGEFB/M/U-deficient mutants (i.e. *gefB-, gefM-* and *gefU-*) in which RasGEFX is expected to be intact, showed a higher frequency of protrusion formation. *gefB-* cells had a highly polarized shape with a shrunken Ras-GTP-enriched domain, but the overexpression of RasGEFB in these cells significantly expanded the area of the protrusion without changing the frequency of the protrusion formation (Figure 5C; Video S5). These observations are consistent with the role of RasGEFB as a spatial regulator in the Ras excitable system. *gefU-* cells showed greatly enhanced protrusion formation due to the increased motility caused by reduced adhesion. RasGEFU overexpression in these cells led to a large Ras-GTP-enriched domain but also less motility (Figures 5C). No obvious changes in the frequency and size of the protrusion formation were observed in RasGEFM KO or OE cells (Figures 5C). Thus, RasGEFX primarily triggers spontaneous Ras excitation to generate a signal for actin-dependent protrusion formation in basic cell motility when no chemoattractant stimulation is present. This spontaneous symmetry breaking in the Ras excitable system leads to anterior-posterior polarity. RasGEFB and RasGEFU modulate the polarity by regulating the sizes of the Ras-GTP-enriched domain and the cell adhesion, respectively.

### Cluster-1 RasGEFs are all dispensable for cAMP-induced Ras activation and chemotaxis

To explore the possible roles of cluster-1 RasGEFs in cAMP signaling and chemotaxis, the response dynamics of Ras activity was examined by applying uniform cAMP stimulations. After stimulation with a sufficiently high concentration (10 μM), a transient membrane translocation of mScarlet-RBD_Raf1_ was observed, peaking at about 5∼7 sec in the WT strain (Figure 6A). Similar responses were observed for *gefB-, gefM-, gefU-* and *gefX-* cells but with slight temporal differences in the cAMP-induced Ras activation (Figure 6B). Dose response curves from 1 pM to 10 μM revealed the mutant cells responded similarly to WT cells (Figure 6C). Therefore, these RasGEFs are unlikely necessary for cAMP-induced Ras activation, suggesting the involvement of other RasGEF clusters in cAMP signaling. Notably, *gefX-* cells were significantly less responsive to cAMP at 1 nM than the other KO strains, suggesting that the spontaneous enhancement of Ras excitability by RasGEFX is essential for a response to low cAMP concentrations.

**Figure 6.**
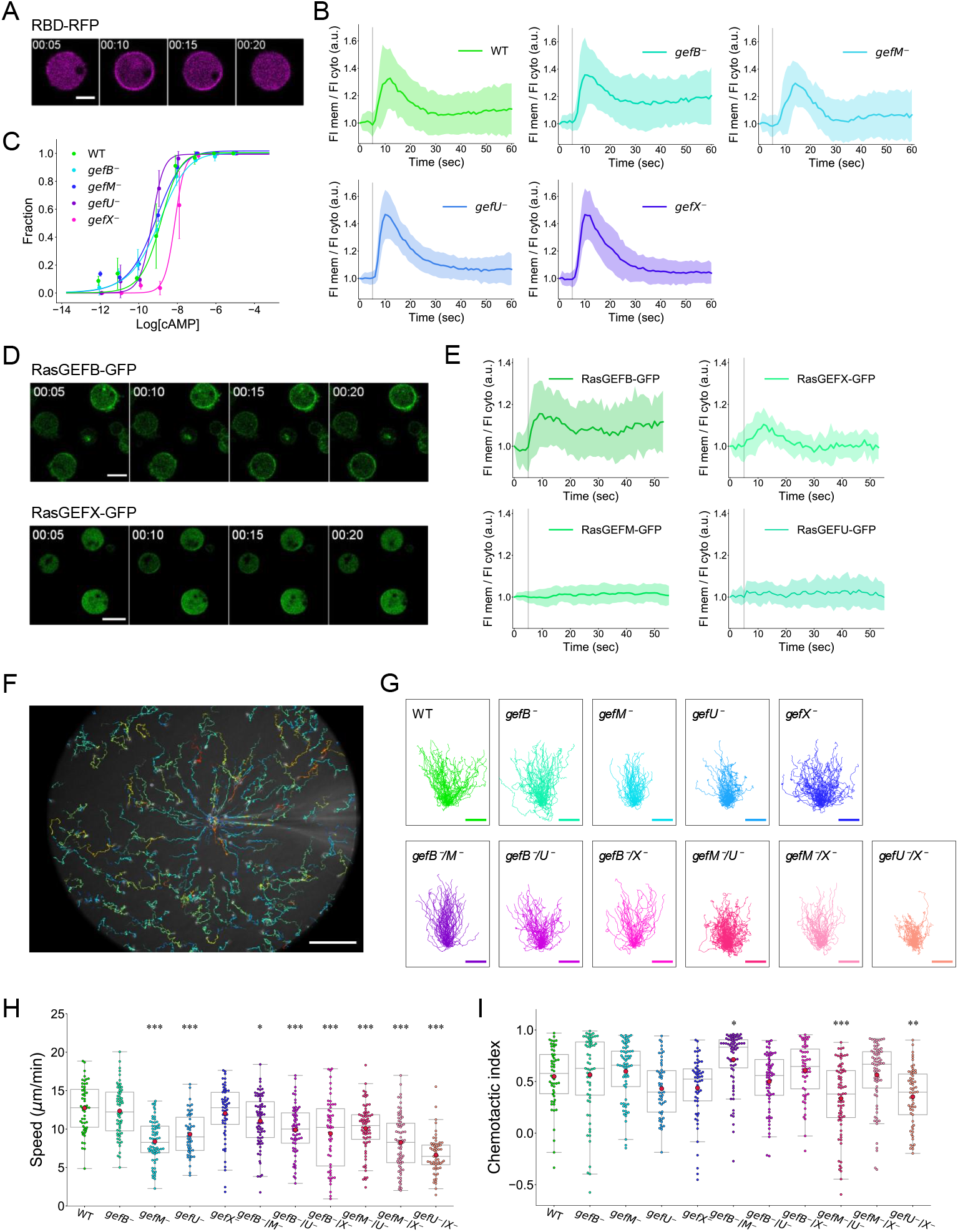
Chemotaxis and cAMP-stimulated Ras activation in RasGEF KO strains. (A) Time lapse images showing the transient membrane localization of RBD_Raf1_-RFP in a wild-type (WT) cell in response to uniform 10 μM cAMP stimulation. Scale bar, 5 μm. Time, min:sec. (B) Quantification of the response in WT (*n* = 67 cells), *gefB-* (*n* = 44 cells), *gefM-* (*n* = 33 cells), *gefU-* (*n* = 50 cells), and *gefX-* (*n* = 56 cells) using mScarlet-RBD_Raf1_ and 10 μM cAMP. (C) Fraction of responsive cells upon stimulation with 1 pM to 10 μM cAMP. *n* > 30 cells from two independent experiments for each concentration. The half-maximum effective concentrations are 1.34 nM (WT), 1.09 nM (*gefB*-), 0.69 nM (*gefM*-), 0.47 nM (*gefU*-) and 7.29 nM (*gefX*-). (D) Time lapse images showing the response of RasGEFB-GFP (*top*) and RasGEFX-GFP (*bottom*) expressed in the respective RasGEF KO strains to 10 μM cAMP. Scale bars, 10 μm. Time, min:sec. (E) Quantification of the response of RasGEFB-GFP (*n* = 47 cells), RasGEFM-GFP (*n* = 20 cells), RasGEFU-GFP (*n* = 24 cells) and RasGEFX-GFP (*n* = 56 cells) using 10 μM cAMP. (F) Trajectories of the cells undergoing chemotaxis in response to 10 μM cAMP for 20 min. Scale bar, 100 μm. (G) Trajectories of WT (*n* = 54 cells), *gefB-* (*n* = 61 cells), *gefM-* (*n* = 70 cells), *gefU-* (*n* = 51 cells), *gefX-* (*n* = 56 cells), *gefB-/M-* (*n* = 70 cells), *gefB-/U-* (*n* = 63 cells), *gefB-/X-* (*n* = 55 cells), *gefM-/U-* (*n* = 72 cells), *gefM-/X-* (*n* = 71 cells) and gefU-/X-(*n* = 61 cells) undergoing chemotaxis in response to 10 μM cAMP for 20 min. Scale bars, 100 μm. (H) Speed of chemotactic migration. (I) Chemotaxis index. (H and I) Closed circles in magenta show mean values. * *p* < 0.05, ** *p* < 0.01, *** *p* < 0.001 as calculated using Dunnett’s test.

Uniform cAMP stimulations induced the transient membrane localization of RasGEFB as well as RasGEFX, albeit to a lesser extent (Figure 6D). RasGEFX-GFP and RasGEFB-GFP responded with a peak at about 5∼7 sec upon stimulation, but neither RasGEFM-GFP nor RasGEFU-GFP showed a peak (Figure 6E). These results indicate that some binding sites for RasGEFX and RasGEFB are produced on the membrane upon cAMP stimulation. One possible candidate for the binding site is active Ras and/or some factors localized in the Ras-GTP-enriched domain, because some mechanisms to recruit RasGEFX and RasGEFB to the Ras-GTP-enriched domain are suggested by the co-localization of these two molecules with Ras-GTP under unstimulated conditions (Figure 4).

Chemotaxis toward cAMP was examined by introducing a glass micropipette filled with 10 μM cAMP among the cells (Figure 6F). Chemotaxis was undisturbed in the KO strains, as seen in the 20-min trajectories of cell migration (Figure 6G; Video S6). The migration speeds of *gefB-* and *gefX-* were comparable with WT, but the speeds of the other mutants were slower. Nevertheless, the speeds for all observed KO types were faster than in the absence of cAMP, showing the chemokinetic effects of cAMP on cluster-1 RasGEF-deficient mutants (Figure 6H; Table S2). The chemotaxis index, a measure of chemotactic accuracy toward the chemoattractant source, showed no significant differences between WT (0.55 ± 0.29) and most mutants. Two exceptions were *gefM-/U-* (0.33 ± 0.35) and *gefU-/X-* (0.35 ± 0.28) (Figure 6I; Table S2). So too was *gefB-/M-*, which showed enhanced chemotaxis efficiency (0.71 ± 0.27). The genetic disruption of RasGEFX, as seen in *gefB-/X-, gefU-/X-* and *gefM-/X-* cells, impaired basic and random cell motility severely without external cAMP (see Figure 3), but their mobilities were recovered as directional motility with cAMP stimulation (Figures 6G-6I). These results indicate that RasGEFB/M/U/X are not necessary for regulating the directionality of cell migration under cAMP gradients and that other RasGEFs may enhance the directional motility instead. Rather, cluster-1 RasGEFs are essential for the basal cell motility that arises independently of chemotactic signaling from environmental guidance cues.

## DISCUSSION

Excitable systems have been identified as key signaling networks for random cell migration as well as chemotactic directional migration in eukaryotic amoeboid cells.^11,49^ They cause Ras-GTP to self-organize, thus generateing the signaling domain on the membrane for the regulation of cell motility.^19,24,29,30^ Herein, by applying a comprehensive imaging analysis of the spatiotemporal dynamics of Ras-GTP in living *Dictyostelium* cells, we found that RasGEFX is primarily required for the spontaneous excitability that generates both the Ras-GTP-enriched domain under physiological conditions and the traveling waves of the domain under conditions with enhanced excitability (Figures 1, 2, and 4). It is also required for basal cell motility (Figure 3). In contrast, RasGEFB/M/U are dispensable for this reaction but work as regulators of the spontaneous dynamics (Figures 1, 2, and 4). They are also required in combination with RasGEFX for random cell migration (Figure 3). RasGEFX and RasGEFB regulate the Ras excitable system with their asymmetric localization on the membrane, while RasGEFU and RasGEFM regulate the excitable system with their uniform localization on the membrane and in the cytosol, respectively (Figure 4). Moreover, RasGEFX and RasGEFB regulate the cytoskeletal dynamics in a different manner (Figure 5), while RasGEFU regulates cell adhesion (Figure 5). Finally, RasGEFB/M/U/X all are dispensable for chemotactic signaling (Figure 6). These findings indicate that a specific set of RasGEFs constitutes spontaneous signal generators for driving the random cell motility that operates independently of external chemotactic signaling.

Excitability has been well documented experimentally in various biological systems, and the characteristic spatiotemporal dynamics has been explained by various theoretical models.^26,49–53^ In general, an excitable system has a threshold for all-or-none excitation, by which spontaneous excitation occurs when the threshold is exceeded by the intrinsic fluctuations of the system itself, and a traveling wave is constantly generated when the system is always over the threshold due to the enhanced excitability.^26,52,53^ A theoretical model for the Ras excitable system predicts that when the RasGEF activity becomes stronger, the spontaneous excitation becomes more frequent and the oscillation periods of the traveling wave become shorter; and when the amount of Ras increases on the membrane, the domain size becomes larger.^24^ The amount of Ras can be regulated by the recruitment and/or the membrane-binding stability during dynamic shuttling between the membrane and cytosol.^54–56^ Based on the different and combinatory roles of RasGEFX and RasGEFB/M/U determined experimentally, we propose a model for the emergence of the spontaneous excitable dynamics of Ras as follows. RasGEFX primarily catalyzes Ras activation on the membrane by its basal activity. This activation triggers the spontaneous excitation leading to symmetry breaking by RasGEFX regulating the temporal properties of the Ras-GTP-enriched domain such as the firing frequency and the oscillation period. RasGEFB/M/U function subsidiarily with respect to the Ras activation. Our model assumes that RasGEFB is involved in the recruitment and/or membrane-binding stability of Ras on the membrane, an effect that can explain its regulation of the spatial properties of the Ras-GTP-enriched domain such as the domain size. RasGEFM and RasGEFU are likely to modulate the spontaneous excitability by their uniform localization in the cytosol and on the membrane, respectively, but their contributions to the spatiotemporal regulation of the Ras-GTP-enriched domain are not so obvious when no guidance cues are present. Because RasGEFM and RasGEFU (GbpD) can be phosphorylated and bound to cGMP under chemotactic stimulation, respectively,^36,44^ they may be important in mediating chemotactic signals to Ras, although they are not essential for cAMP-induced Ras activation.

A theoretical consideration of this excitable system suggests positive feedback mechanisms that amplify signals for the all-or-none response. The co-localization of RasGEFB and RasGEFX with the Ras-GTP-enriched domain on the membrane in an asymmetric manner suggests that the recruitment of RasGEFB and RasGEFX to the membrane depends on Ras-GTP and/or some factors localized in the Ras-GTP-enriched domain (Figure 4), further suggesting positive feedback between RasGEFX/B and Ras-GTP. RasGEFX has a primary structure similar with that of mammalian SOS (Son of Sevenless) in the Ras/MAP kinase pathway (Figure S3E).^57,58^ Extensive structural and biochemical analyses of SOS have revealed positive-feedback loops, in which SOS can bind to Ras-GTP and modify its basal GDP/GTP exchange rate for Ras-GDP.^55,59–61^ This mechanism can constitute an amplifying mechanism for the burst of Ras activation locally on the membrane, an effect favorable for Ras-GTP-enriched domain formation. RasGEFB has a common RasGEF domain but no other recognizable signaling domains.^34^ More studies are needed to understand the positive-feedback mechanisms for the excitable systems in eukaryotic chemotaxis, especially for the spontaneous generation of the Ras-GTP-enriched domain.

RasGEFX and RasGEFB regulate temporally and spatially the Ras excitability and cytoskeletal dynamics, leading to the regulation of the frequency of the protrusion formation and size of the protrusion, respectively. Both RasGEFX and RasGEFB co-localized with the Ras-GTP-enriched domain at lamellipodia-like protrusions and macropinocytic cups; thus, the spatiotemporal dynamics of actin cytoskeleton-dependent protrusions is diverse depending on the balance of the expression levels of these two RasGEFs. For example, RasGEFB can enlarge the Ras-GTP-enriched domains when overexpressed to cause a modal transition of the cell motility from amoeboid migration to keratocyte-like gliding with a fan-shape (Video S5). Such a transition in the motility mode has been well documented with changes of the spatiotemporal dynamics of the excitable system in *Dictyostelium* cells.^62^ In addition, the overexpression of RasGEFX promoted micropinocytosis (Figure 5), but no such promotion was observed for the overexpression of RasGEFB, suggesting their different roles in the regulation of the cytoskeletal dynamics. Because RasGEFX and RasGEFB share a common RasGEF domain but differ in other regions, including Ser/Thr protein kinase for RasGEFX (Figure S3E), they may share regulatory functions for Ras excitability but also have other signaling functions for cell motility. As reported previously, RasGEFU is involved in Rap1 activation and thus regulates cell adhesion.^36^ Given that spontaneous cell migration requires concerted regulation of the actin cytoskeleton and adhesion, RasGEFB/U/X may contribute to the coordination by their common function in the excitable Ras system and their differential function in the motile system.

Our observation that RasGEFB/M/U/X are not required for cAMP-induced Ras activation or chemotaxis may be explained at least partly by the network structure for chemotaxis signaling in *Dictyostelium discoideum*. At least 14 genes for Ras/Rap small GTPases have been found in the *Dictyostelium* genome.^33^ Among them, at least 4 Ras/Rap GTPases are involved in chemotactic signaling, in which four parallel signaling pathways, including the RasG/RasD/PI3K, RasC/TorC2, Rap1/sGC and PLA2 pathways, mediate chemotactic signals from cAMP receptors to the actin cytoskeleton for cell motility.^11,31^ Because RBD_Raf1_-RFP and mScarlet-RBD_Raf1_ primarily detect active RasG,^41^ our results indicate that each RasGEFB/M/U/X regulates at least the RasG/PI3K pathway directly or indirectly for spontaneous signal generation. However, even if each RasGEFB/M/U/X mediates chemotactic signals to RasG, the loss of their function only causes defects in the RasG/PI3K pathway. Consistently, rasGcells can exhibit chemotaxis with only slight defects under cAMP gradients,^63^ in which the three signaling pathways besides the RasG/PI3K pathway mediate the chemotactic signals. Furthermore, RasGEFA/G/K/R/T in clusters 4 and 5 are phosphorylated by chemoattractant stimulation,^44^ and RasGEFA/C/H/F/R/Q/U are involved in Ras/Rap activation upon cAMP stimulation.^37,39,40^ Future studies should seek to identify the RasGEFs responsible for the chemoattractant-induced activation of the Ras excitable system by using a systematic and comprehensive screening method.

## ACKNOWLEDGEMENTS

We thank Dr. Peter Karagiannis for critical reading of the manuscript. This work was supported by funds from the Japan Society for the Promotion of Science (JSPS) Grant-in-Aid for JSPS Research Fellow grant no. 22KJ2207 to KI, JSPS KAKENHI grants no. 19H00982 to MU and no. 19H05798 to SM, Japan Science and Technology Agency grant no. JPMJPR1879 to SM and JPMJCR21E1 to MU, and Japan Agency for Medical Research and Development grant no. JP20gm0910001 to MU.

## AUTHOR CONTRIBUTIONS

Conceptualization, K.I., S.M., and M.U.; Methodology, K.I., S.M., and M.U.; Software, K.I.; Validation, K.I., S.M., and M.U.; Formal analysis, K.I.; Investigation, K.I.; Resources, K.I., S.M., and M.U.; Data curation, K.I.; Writing -original draft, K.I.; Writing -review &editing, K.I., S.M., and M.U.; Visualization, K.I.; Supervision, M.U.; Project administration, S.M., and M.U.; Funding acquisition, K.I., S.M., and M.U.

## MATERIAL AND METHODS

### Cell strains

*Dictyostelium discoideum* wild-type Ax2 was used as the parental strain. All cell lines were grown in HL5 medium (Formedium, UK) supplemented with penicillin and streptomycin, 100 ng/mL folic acid and 5 ng/mL vitamin B12 at 21°C.^64^ The strains overexpressing one of 25 RasGEFs in GFP-tagged form were generated by the electroporation of extrachromosomal plasmids for stable expression into the parental strain cells. The plasmids were generated by cloning RasGEF genes, which were amplified from wild-type genomic DNA by PCR using the primers listed in Table S2, into the BglII site of pHK12neo. The transformants were selected and maintained under 20 μg/mL G418. The strains co-overexpressing RBD_Raf1_-RFP or mScarlet-RBD_Raf1_ and RasGEF-GFP were generated by the electroporation of extrachromosomal plasmids for the stable expression of the Ras-GTP probe into RasGEF-GFP OE strains. The generation of the plasmid for RBD_Raf1_-RFP is described elsewhere.^24^ The plasmid for mScarlet-RBD_Raf1_ pDM1208 was provided by National Bio Resource Project (NBRP). The transformants were selected and maintained under 40 μg/mL Hygromycin B and 20 μg/mL G418. The expression of RBD_Raf1_-RFP and mScarlet-RBD_Raf1_ was unstable, and the cells were observed microscopically within about 15 days of the electroporation. The single gene knockout strains, *gefX-, gefB-, gefU-* and *gefM-*, were generated using CRISPR/Cas9 by introducing extrachromosomal plasmids for the transient expression of both sgRNA and Cas9-NLS-GFP into the parental strain cells.^65^ The plasmids were generated by cloning the target-site dsDNA fragments (Table S2) into BpiI sites located between the isoleucine tRNA promoter and tracrRNA sequence in pTM1285 by Golden Gate assembly.^66^ The transformants were selected under 20 μg/mL G418 for 1 day and then maintained in the absence of G418. For cell cloning, we plated the cells on 5LP agar plates (Lactose 5.0 g, Bacto Peptone 5.0 g, Bacto agar 15.0 g) with Escherichia coli B/r and incubated them for 3–4 days until plaque formation. We transferred the individual plaques to 96-well plates (Thermo Fisher) containing HL5 in the absence of G418. DKO strains, including *gefB-/M-, gefB-/X-* and *gefM-/X-*, were generated by introducing the same CRISPR/Cas9 plasmids targeting *gefM, gefX* and *gefM* as described above into the strains of *gefB*-, *gefB*- and *gefX*-, respectively. The strains, including *gefB-/U-, gefM-/U-* and *gefU-/X-*, were generated by homologous recombination targeting *gefU*.^67^ Two fragments of RasGEFU gene, from bases 867 to 1641 and 2106 to 2993, were amplified by PCR using genomic DNA as a template and the primers listed in Table S2, and the blasticidin S resistance (BSR) gene was inserted between them by fusion PCR. The fusion PCR was performed in 10 μL total volume using 1 ng of two fragments of RasGEFU gene, the fragment of BSR gene and 5 pmol of primerA and primerD (Table S2). The fusion PCR conditions were as follows: heating to 94°C for 2 min; 30 cycles of 94°C for 30 sec, 50°C for 20 sec and 68°C for 4 min; followed by a final extension for 4 min. The transformants were selected under 10 μg/mL blasticidin S. We performed cloning with CRISPR-Cas9.

### Electroporation of *Dictyostelium* cells

The cultured cells were collected by centrifugation at 500×g for 2 min and resuspended at 1×10^7^ cells/mL in H50 buffer (50 mM KCl, 20 mM HEPES, 10 mM NaCl, 5 mM NaHCO_3_, 1 mM NaH_2_PO_4_, 1 mM MgSO_4_, pH 7.0). A 100 μL cell suspension was mixed with 5 μg plasmid and incubated on ice for 5 min. The cell suspension was transferred to a cuvette with a 1-mm gap and set in an electroporator (BTX) to give a shock of 500 μV for 100 μsec repeated 2 times at an interval of 5 sec. The cell suspension was kept on ice for 2 min and transferred to a culture dish. After a 2-min incubation at 21°C, 10 mL of HL5 medium was added to grow the cells at 21°C. The drugs used for the selection were added 12∼24 hours after the electroporation.

### Confirmation of gene knockout

Genomic DNA was isolated from the transformants, and the nucleotide sequence at the target site was confirmed as follows. Cells were collected from plaque on 5LP agar plates and were resuspended in lysis buffer (200 mM Tris-HCL pH8.4, 500 mM KCl, 1.75 μM MgCl_2_, 0.5% NP40, 20 μg/mL proteinase K and 20 μg/mL RNase), and the suspension was incubated at 56°C for 50 min and 95°C for 10 min to inactivate proteinase K. A DNA fragment encompassing the target site was amplified by PCR using the lysate as a template, the primers listed in Table S2, and SapphireAmp Fast PCR Master Mix (Takara). The PCR conditions were as follows: heating to 94°C for 2 min; 35 cycles of 94°C for 30 sec, 50°C for 45 sec, and 68°C for 1 min/kbp; followed by a final extension for 5 min. The amplified fragment was subjected to gel electrophoresis, where the band shift due to the insertion of the BSR cassette was confirmed in the case of homologous recombination and purified using a PCR purification kit (NEB), in which the nucleotide sequence was determined (Genewiz). We confirmed that the deletion or insertion of some bases by CRISPR/Cas9 led to an insertion of the stop codon in the coding sequence, in which the nucleotide sequence was determined (Genewiz) using the screening primers listed in Table S2.

### Cell preparation for microscopy

Cultured cells were starved as follows. The cells were washed three times with development buffer without Ca^2+^ and Mg^2+^ (DB−; 5 mM Na_2_HPO_4_ and 5 mM KH_2_PO_4_, pH 6.5) by centrifugation at 500 × g for 2 min. The cells were resuspended at 3.0×10^6^ cells/mL in DB+ (5 mM Na_2_HPO_4_, 5 mM KH_2_PO_4_, 2 mM MgSO_4_, 0.2 mM CaCl_2_, pH 6.5), and 1 mL cell suspension was plated on a 35-mm dish (IWAKI). To observe subcellular localization and spontaneous motility, the cells were incubated for 3 h at 21°C. To observe chemotaxis, the cells were incubated for 5 h at 21°C. After the starvation period, the cells were washed twice with DB- and resuspended at 3.0×10^6^ cells/mL in DB+ on ice until use.

### Fluorescence microscopy

Laser scanning confocal microscopy (A1, Nikon) was performed on the starved cells described above. RFP and mScarlet tagged to RBD_Raf1_ were excited with a 561 nm laser (Coherent), and 570-620 nm fluorescence light was detected by a detector (A1-DU4, Nikon) equipped with the microscope. GFP tagged to RasGEFs was excited with a 488 laser (Coherent), and 500-550 nm light was detected by the detector. An emitted fluorescence was imaged with a 60x objective lens (CFI Apo TIRF 60X Oil, NA 1.49, Nikon). To observe Ras-GTP and RasGEF dynamics in the presence of caffeine (Figures 1, 2, and 4), 150 μL cell suspension was mixed with 150 μL DB+ containing 8 mM caffeine (Wako) and 20 μM latrunculin A (Sigma-Aldrich) and placed on a 35-mm glass bottom dish (12-mm glass in diameter; Iwaki). After the cells were allowed to adhere to the glass surface for 20 min, the images were acquired at a time interval of 5 sec for 20 min.^13^ To investigate the relationship between RasGEF expression levels and Ras-GTP dynamics (Figure 4), a snapshot image of RasGEF-GFP was acquired at the beginning of the image acquisition. To observe Ras-GTP dynamics in the absence of caffeine (Figure 2), a 50 μL cell suspension was mixed with 100 μL DB+ and 150 μL DB+ containing 20 μM latrunculin A and placed on a 35-mm glass bottom dish, in which the cell density was reduced to 1/3 of the sample containing caffeine to suppress intercellular communication via secreted cAMP. After the cells were allowed to adhere to the glass surface for 20 min, the images were acquired every 5 sec for 20 min. To observe Ras-GTP and RasGEF simultaneously in migrating cells (Figure 5), a 50 μL cell suspension was mixed with 250 μL DB+, placed on a 35-mm glass bottom dish, and the cells were allowed to adhere to the glass surface for 20 min.^13^ The images were acquired every 5 sec for 5 min. To observe the response to cAMP (Figure 6), the cell suspension was mixed with DB+ containing latrunculin A at a final concentration of 10 μM, and a 20 μL cell suspension was placed on a 35-mm glass bottom dish. The cells were allowed to adhere to the glass surface for 20 min. The images were acquired every 1 sec for 1 min. When 5 sec passed after starting the image acquisition, 180 μL DB+ containing cAMP was added at a final concentration of 1 pM-10 μM.

### Cell migration imaging

Starved cells were washed twice with DB- and resuspended at 5.0×10^4^ cells/mL in DB+. 300 μL cell suspension was placed on a coverslip (MATSUNAMI) that was first washed by sonication in 0.1 N KOH for 30 min and then 99.5% ethanol. The cells were allowed to adhere to the glass surface for 20 min. 1 mL DB+ was added, and the cells were allowed to settle for another 20 min. The cells were observed under an inverted microscope (IX71, Olympus) with a 20X objective lens (LCACHN20XPH, NA 0.4, Olympus). Images were acquired with a time-lapse camera (DS-2MBW, Nikon) every 5 sec for 30 min. To examine chemotaxis, the concentration gradient of cAMP was generated using FemtoJet (Eppendorf): DB+ containing 10 μM cAMP was filled in a glass micropipette (Femtotips, Eppendorf) and released by applying pressure at 50 hPa.

### Kymograph of the fluorescence intensity on the cell membrane

The spatiotemporal dynamics of RBD_Raf1_-RFP, mScarlet-RBD_Raf1_ and RasGEF-GFP in each cell were analyzed using a kymograph obtained as follows. The fluorescence intensity on the cell membrane was measured in space along the circumference of the cell divided into 90 sections of 4 degrees each and in time over 241 frames in the movie. This measurement yields a two-dimensional matrix data for the *i*-th cell (*i* = 1,2,3, … *N*) written as *mFI*_*i*_(*s*Δ*θ, f*Δ*t*), where *s* (*s* = 1,2,3, … 90) and *f* (*f* = 0,1,2, … 240) denote the section and frame numbers with the intervals Δ*θ* = 4 degrees and Δ*t* = 5 sec. The data were displayed on the horizontal and vertical axes as a kymograph of 90 pixels × 241 pixels.

### Quantification of domain size and period

The domain size was determined by binarizing the kymograph with the method of Li. The following process was performed to remove the influence of noise. In a binary image, each pixel was selected along with the immediately 5 adjacent left and right pixels were extracted (total of 11). If the majority of these pixels had a pixel value of 255, then the selected pixel was set to 255. Conversely, if the majority of these pixels had a pixel a value of 0, then the selected pixel was set to 0. From the obtained binary images, the angle of the domain was calculated at each time point, and the angles were then averaged (Figure S2A). For the period, a temporal autocorrelation function of the fluorescence intensity at an arbitrary section of the cell circumference was calculated as,

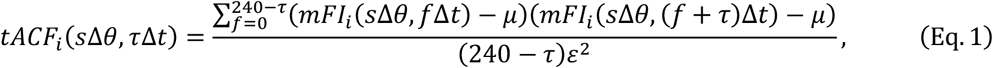

where *μ* and *ε* denote the mean and SD of the fluorescence intensity, respectively. *tACF*_*i*_ was averaged over 90 sections, in which the lag-time, *τ*Δ*t*, giving the first peak was quantified as the period.

### Classification of Ras-GTP dynamics into four patterns

The spatiotemporal dynamics of Ras-GTP that spontaneously emerged in individual cells of each OE/KO strain were classified into four patterns: no domain, transient domain, traveling wave and uniform localization, by hierarchical clustering.^68^ The spatiotemporal dynamics in each cell was represented by a one-dimensional vector individually calculated from the kymograph as described below and subjected to hierarchical clustering using scipy 1.2.1. cluster.hierarchy. An iterative calculation was performed as follows. The initial condition of a single cell was represented by a component of the ensemble to be clustered, in which the centroid was calculated from the kymograph using the one-dimensional vector described below. The distance between two components was calculated for all possible pairs, and those with the nearest distance were combined as a new component, in which the centroid was calculated by averaging the two centroids. The calculation was repeated until all components were combined into one. The distance was calculated by Ward’s method written as,

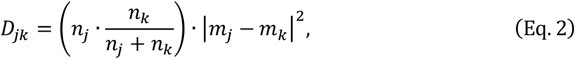

where *n*_*j/k*_ and *m*_*j/k*_ denote the number of combined components and the centroid of the *j*/*k*-th component, respectively. |*m*_*j*_ − *m*_*k*_|^2^ is the squared Euclidean distance between the centroids. Each pattern’s clusters were determined based on the dendrogram by setting a threshold (Figure S1).

First, the cells were classified into two subgroups, those exhibiting a traveling wave and those exhibiting other patterns (Figures S1A and S1B). For this purpose, the following three quantitative expressions of the kymograph were exploited for the clustering to reveal periodicity in the dynamics: (1) The kymograph was normalized by the maximum fluorescence intensity around the cell periphery at each time point and smoothed with a Gaussian filter of *σ* = 1, which is described as a vector, 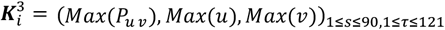; (2) A temporal autocorrelation function for up to *τ* = 120 sec, which is the duration when the periodicity is remarkable, was calculated from the kymograph smoothed with a Gaussian filter of *σ* = 5, which was described as a vector, 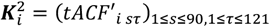; and (3) A power spectrum was obtained by Fourier transforming 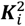 using,

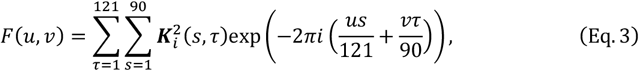

which was written as,

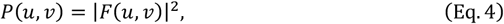

where (*u, v*) is the coordinates in the frequency domain. The maximum value, *Max*(*P*(*u, v*)), and its coordinates, *Max*(*u, v*), of the power spectrum were described as a vector, 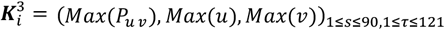. 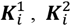 and 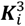 were combined and linearly transformed into a one-dimensional vector, which was subjected to the hierarchical clustering.

The cells exhibiting other patterns were further classified into two subgroups, those exhibiting transient domains and those without domains (Figures S1C and S1D). For this purpose, the following two quantitative expressions of the kymograph were exploited for the clustering to reveal a spatial heterogeneity in the dynamics: (4) The kymograph was normalized as mentioned above, as a vector, 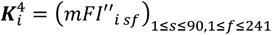; and (5) A spatial autocorrelation function was calculated for up to *τ* = 240 sec from the kymograph smoothed with a Gaussian filter of *σ* = 1 as a vector, 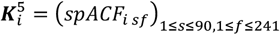, written as,

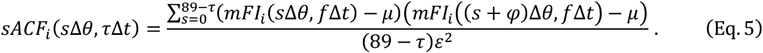

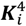 and 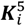 were combined and linearly transformed into a one-dimensional matrix, which was subjected to hierarchical clustering.

Next, the cells with other patterns were classified into two subgroups, those exhibiting a uniform domain and those exhibiting no domain (Figures S1E and S1F). For this purpose, the following quantitative expression of the kymograph was exploited for the clustering to reveal the fluorescence intensity on the plasma membrane. The fluorescence intensity of the plasma membrane was calculated by the spatio-temporal average of the kymograph: ⟨*mFI*_*i*_(*s*Δ*θ, f*Δ*t*)⟩; and the fluorescence intensity of the of the cytoplasm was calculated by the temporal average of the kymograph written as ⟨ *cFI*_*i*_(*f*Δ*t*)⟩. The ratio of the fluorescence intensities at the plasma membrane and the cytoplasm was described as,

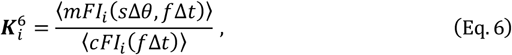

and 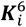 was subjected to hierarchical clustering.

### Classification of RasGEFs into Clusters

The statistical distributions of the domain size and period of each RasGEF OE strain were displayed as a single heat map in domain size-period space (Figure S2A). The widths of the bins were determined based on Freedman-Diaconis’ rule to be 30 sec and 20 degrees assuming the cells showed traveling waves and domains, respectively. The data of all cells irrespective of the domain pattern were incorporated into the heat map by setting the summed probability density in each pattern equal to the fraction. Based on the extent of the excitability, the period of the cells exhibiting no domain (domain size = 0 degrees) was approximated as 460 sec, or the maximum value of the heat map, and that of the cells exhibiting uniform localization (domain size = 360 degrees) was approximated as 0 sec, or the minimum value. The period of cells exhibiting a transient domain of various sizes was approximated as 460 sec. This approximation was based on a previous simulation that showed the period becomes longer as the excitability decreases^24^ and the fact that less than 1% of cells showing a traveling wave had a period of 450 sec or longer. These heat maps were used for hierarchical clustering with the ward method after linearly transforming them into one-dimensional vectors to classify RasGEFs based on the spatiotemporal characteristics of the Ras-GTP dynamics.

### Subcellular localization analysis of RasGEFs

Time-lapse movies of RasGEF-GFP-expressing cells in the presence of latrunculin A acquired by confocal microscopy at 5-sec intervals for 5 min were used for the analysis. After the effect of fluorescence photo-bleaching was corrected based on an approximation with an exponential function, a maximum projection of the movie stack was performed to obtain a still image to exclude artifacts of transient and localized reductions in the fluorescence intensity caused by intracellular vesicles that do not contain GFP. An image of the cell was elliptically approximated and divided into 10 regions by concentric ellipsoids at equal intervals: region 1 to region 10 from the inside to the outside of the cell (Figure S4). The fluorescence intensity was measured in the *h*-th region, 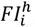, and all regions, 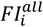, in the *i*-th cell. An index of localization to the *h*-th region, 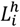, was written as,

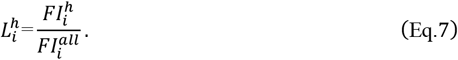

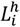 was averaged over *N* cells (*N* ≥ 8). For uniform localization it approximates 1, and for membrane localization it is less than 1 in the inner regions and greater than 1 in the outer regions.

### Co-localization analysis between RasGEFs and Ras-GTP

To examine the co-localization of RasGEFs and Ras-GTP, a cross-correlation function between the fluorescence intensity time series of GFP tagged to RasGEF and RFP or mCherry tagged to RBD, 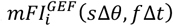 and 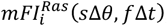, respectively, simultaneously observed on the cell membrane in the *i*-th cell was calculated as,

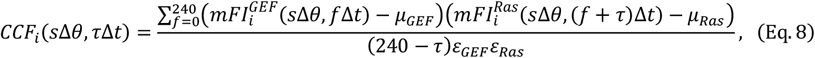

where *μ*_GEF/Ras_ and *ε*_GEF/Ras_ denote the mean and SD of the fluorescence intensities, respectively. *CCF*_*i*_ was averaged over 90 sections, and the coincidence between two traveling waves was confirmed.

### Analysis of cytoskeleton-dependent protrusion dynamics

Pseudopod formation frequency and pseudopod size were quantified in phase-contrast microscopy images of RasGEF KO cells and laser confocal scanning microscopy images of RasGEF-GFP OE cells (Figure 5B). In an image of a cell at an arbitrary time, the pseudopodial region was determined as the region remaining after the subtraction of an image of the same cell acquired 10 sec before. Here, lamellipodia-like protrusions and macropinocytosis-associated protrusions were not distinguished. If a pseudopod continued to extend in the same direction, it was considered a single pseudopod. If a pseudopod formed from a different region, it was considered a newly formed pseudopod. The number of newly formed pseudopods during 5 min was counted, and the pseudopod formation frequency was calculated as the number per minute. Pseudopod size was calculated as the time average ratio in length of the perimeter section along the pseudopodial region to the entire perimeter of the cell preceding the image subtraction.

### Cell migration analysis

The *x*- and *y*-coordinates of the centroid of the cell were obtained using laboratory-made software.^5,6^ The migration trajectory was obtained at 5-sec intervals (Δ*t* = 5 sec) for 30 min in a spontaneous motility assay (*f*Δ*t* = 30 min; *f* = 1, …, 360) and 20 min in a chemotaxis assay (*f*Δ*t* = 20 min; *f* = 1, …, 240). The migration speed, *v*_*i*_, of the *i*-th cell (*i* = 1, …, *N*) was quantified by averaging the instantaneous speed measured in a unit time window of *t*′Δ*t* = 120 sec over the trajectory as

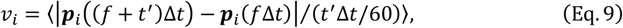

where ***p***_*i*_(*t*)denotes the position at time *t*. The mean squared displacement (MSD) is defined as

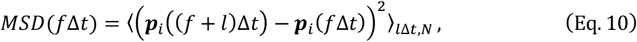

where *l* is the frame number of the lag time. The chemotaxis index, *CI*, of a cell, which indicates the efficiency of the chemotaxis, was quantified by averaging the cosine of the angle between two vectors, 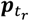 and ***p***_*c*_, which are the direction vectors from the start to the end position during an arbitrary 120 sec and from the initial position, ***p***_*initial*_, to the micropipette position, respectively, over the trajectory:

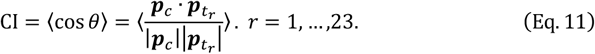

### cAMP response analysis

The fluorescence intensity of RBD_Raf1_-RFP was measured and averaged across the cell membrane, divided by the fluorescence intensity averaged in the cytoplasm, and normalized by the mean value before stimulating individual cells. A maximum intensity was detected in the time series after the moving average was taken and compared to a threshold value to evaluate the response of RBD, which showed transient translocation to the cell membrane. Cells that exhibited a larger maximum intensity than the threshold were regarded as responsive, and a fraction of these cells was examined across the cAMP concentrations 1 pM to 10 μM. The concentration which induced a response in 50% of the cells (i.e., EC_50_) was estimated by fitting the dose-response plot to the equation,

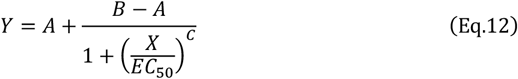

where *A, B, c, X* and *Y* denote the minimum value, the maximum value, Hill coefficient, cAMP concentration, and the fraction of responsive cells.

## SUPPLEMENTAL INFORMATION

### Supplemental Tables

**Table S1. Statistical characteristics of spontaneous Ras-GTP dynamics and cell motility**

**Table S2. List of primers used in this study**

### Supplemental video legends

**Video S1**. Spontaneous dynamics of Ras-GTP detected with RBD_Raf1_-RFP. Four patterns of Ras-GTP membrane localization in wild-type cells (*top row*). Representative cells of RasGEF OE strains (*bottom four rows*). The cells were treated with caffeine and latrunculin A. Scale bars, 5 μm (*white*) and 10 μm (*magenta*). Time, min:sec.

**Video S2**. Spontaneous dynamics of Ras-GTP in cluster-1 RasGEF KO (*top four rows*) and OE (*bottom*) cells in the presence (*top two rows*) or absence (*bottom three rows*) of caffeine with latrunculin A treatment. Scale bar, 10 μm. Time, min:sec.

**Video S3**. Spontaneous migration of cluster-1 RasGEF KO and OE cells. Scale bar, 100 μm. Time, min:sec.

**Video S4**. Spontaneous dynamics of RasGEF (GFP) and Ras-GTP (RFP or mScarlet) simultaneously observed in the respective RasGEF KO cells treated with latrunculin A and caffeine. Scale bars, 5 μm. Time, min:sec.

**Video S5**. Spontaneous dynamics of RasGEF (GFP) and Ras-GTP (RFP or mScarlet) simultaneously observed in the respective RasGEF KO cells without latrunculin A and caffeine. Bottom two rows show Ras-GTP observed in the RasGEF KO cells with different expression levels of RasGEF. Scale bar, 10 μm. Time, min:sec.

**Video S6**. Chemotaxis of cluster-1 RasGEF KO cells in response to 10 μM cAMP. Scale bar, 100 μm. Time, min:sec.

---

**Figure S1.**
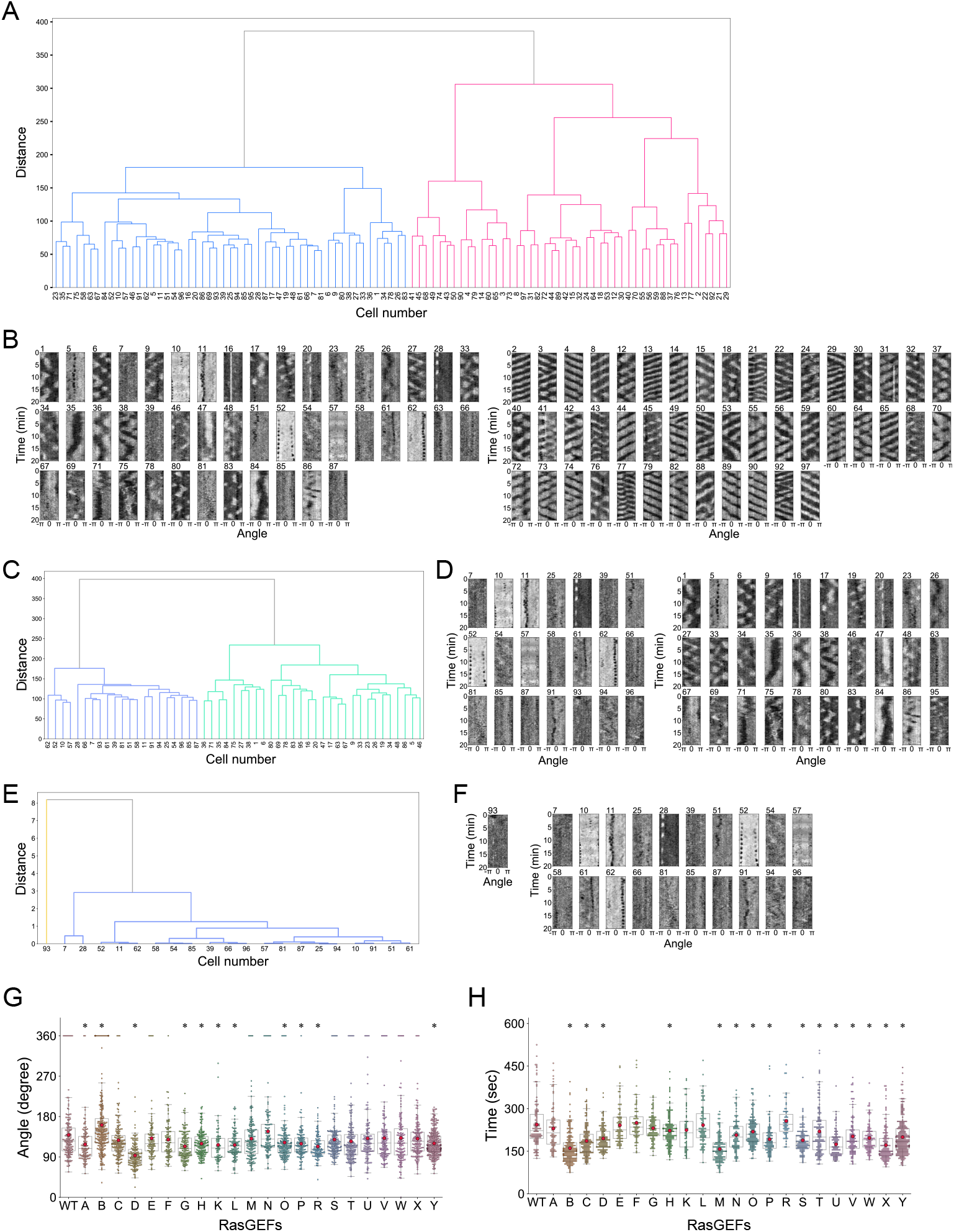
Classification of domain patterns by hierarchical clustering. (A) Hierarchical clustering to classify wild-type cells into two subgroups: those exhibiting traveling waves (*pink*) and those exhibiting other patterns (*blue*). (B) Kymographs obtained from the cells classified in (A) (*right*, traveling waves; *left*, other patterns). (C) Hierarchical clustering to classify cells exhibiting other patterns into two subgroups: those exhibiting transient domains (*green*) and those without domains (*blue*). (D) Kymographs obtained from the cells classified in (C) (*right*, transient domains; *left*, no domains or uniform localization). (E) Hierarchical clustering to classify the cells without domains into two subgroups, those exhibiting uniform localization (*yellow*) and those exhibiting no domain (*blue*). (F) Kymographs obtained from the cells classified in (E) (*left*, uniform localization; *right*, no domain). (G and H) Domain size (G) and period (H) in individual cells of RasGEF OE strains. Closed circles in red indicate mean values, and horizontal lines in box plots indicate median values. * *p* < 0.05 using Dunnett’s test. The numbers of cells analyzed are summarized in Table S1.

**Figure S2.**
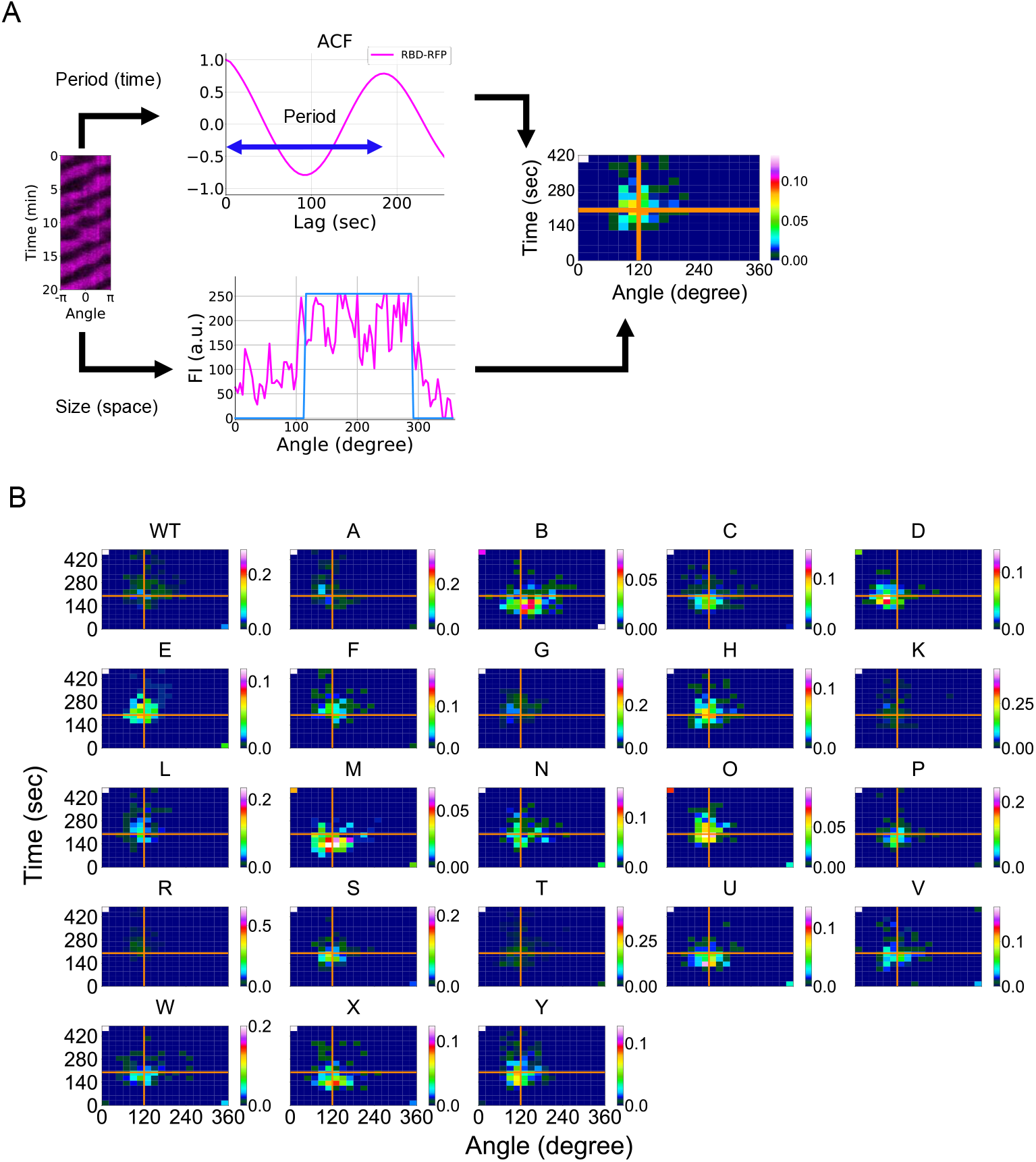
Heat map showing spatiotemporal characteristics of Ras-GTP. (A) Schematic showing the generation of a heat map representing the statistical distribution of the period and domain size in individual cells. Orange lines indicate the mean values of the wild-type strain. (B) Heat maps obtained from the RasGEF OE strains. The number of cells analyzed is summarized in Table S1.

**Figure S3.**
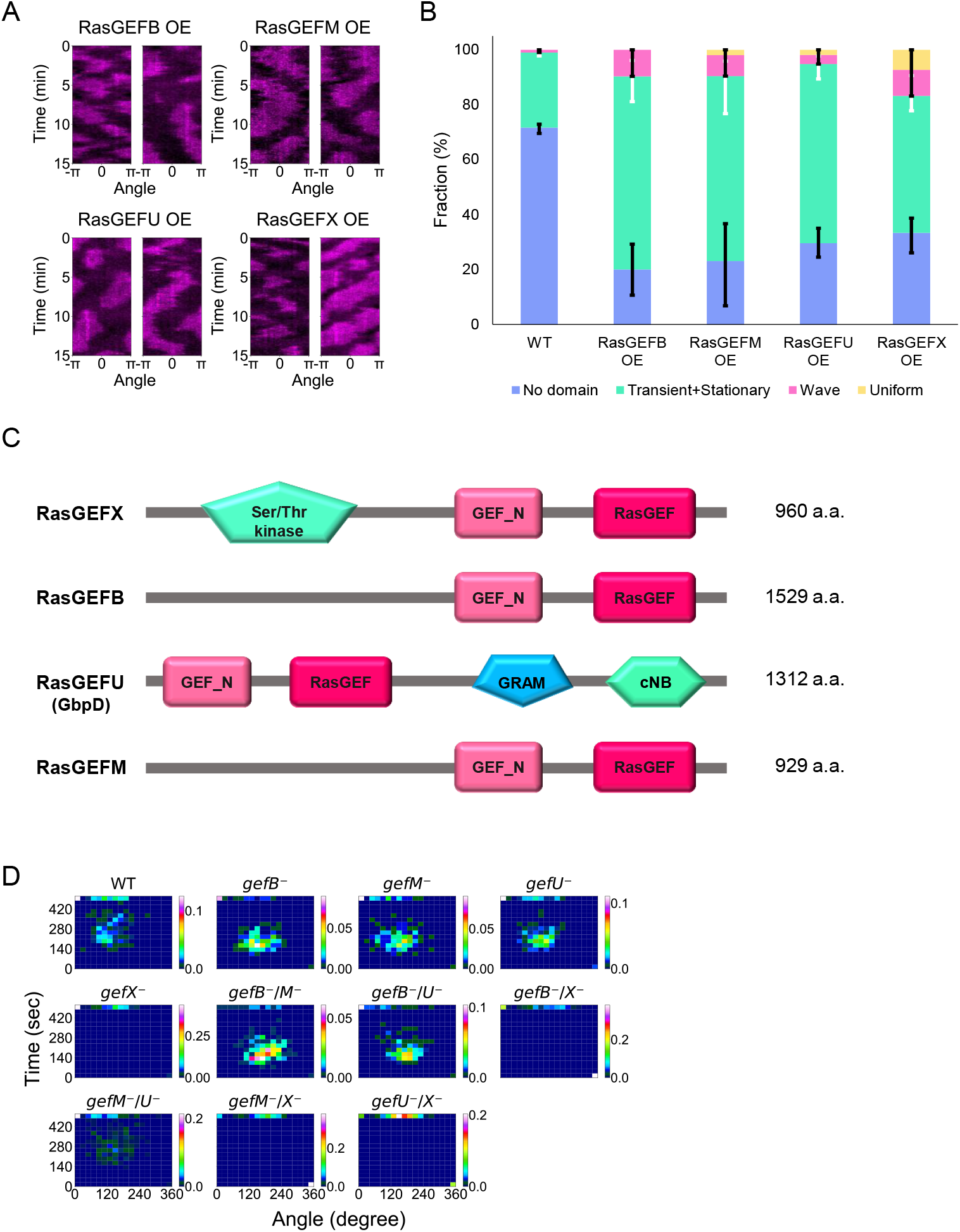
RasGEFX is required for traveling waves of Ras-GTP. (A) Representative kymographs of RBD_Raf1_-RFP in the RasGEF OE strains without caffeine. (B) The fraction of cells showing each domain pattern: no domain (*blue*), non-oscillatory domains (*green*), traveling wave (*magenta*), uniform (*yellow*). The means and SDs of 2 or 3 independent experiments are shown (WT, *n* = 91 cells; RasGEFB OE, *n* = 83 cells; RasGEFM OE, *n* = 52 cells; RasGEFU OE, *n* = 58 cells; RasGEFX OE, *n* = 84 cells). (C) Primary structures of RasGEFX, RasGEFB, RasGEFU, and RasGEFM. (D) Heat maps representing the statistical distribution of the period and domain size in RasGEF KO strains.

**Figure S4.**
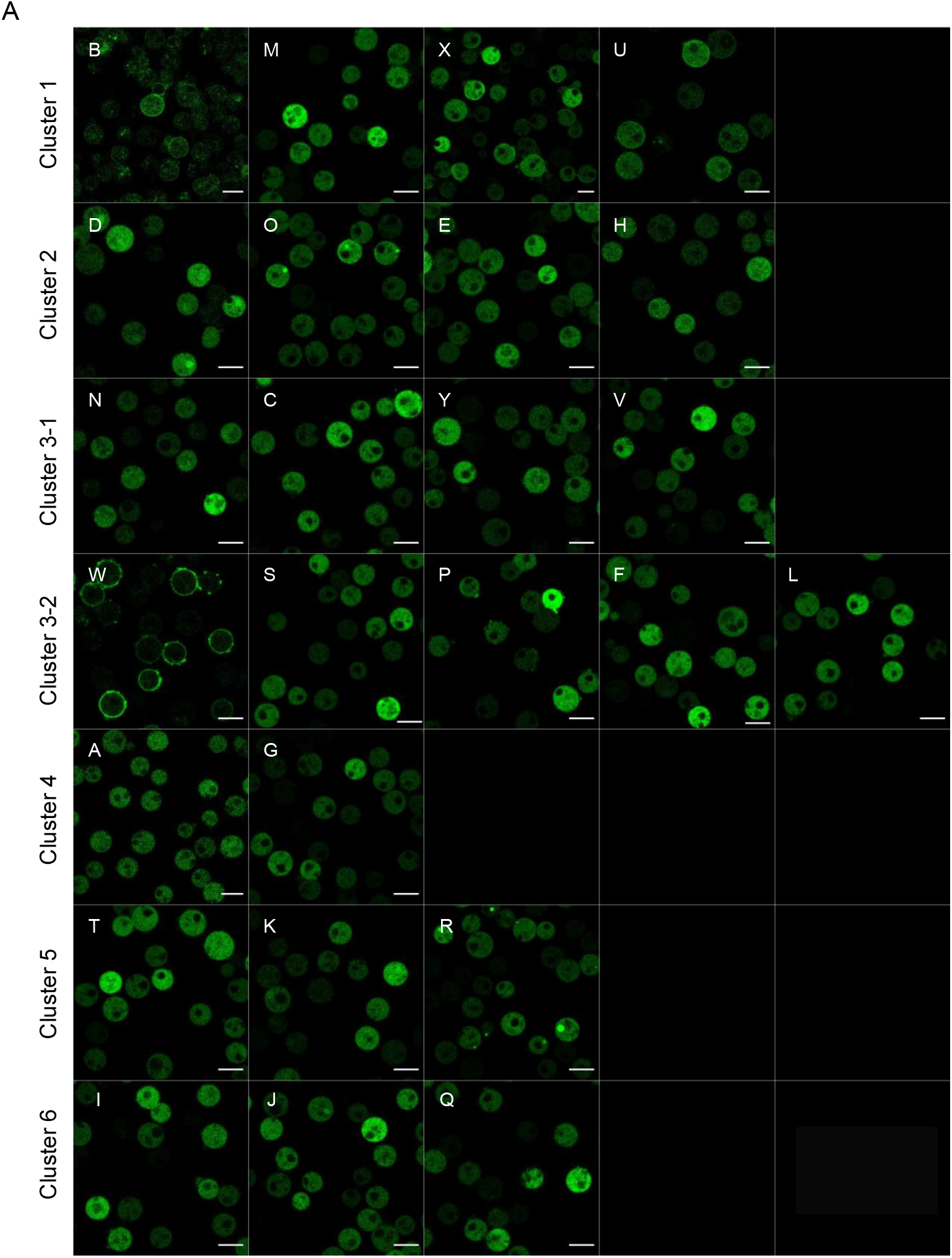
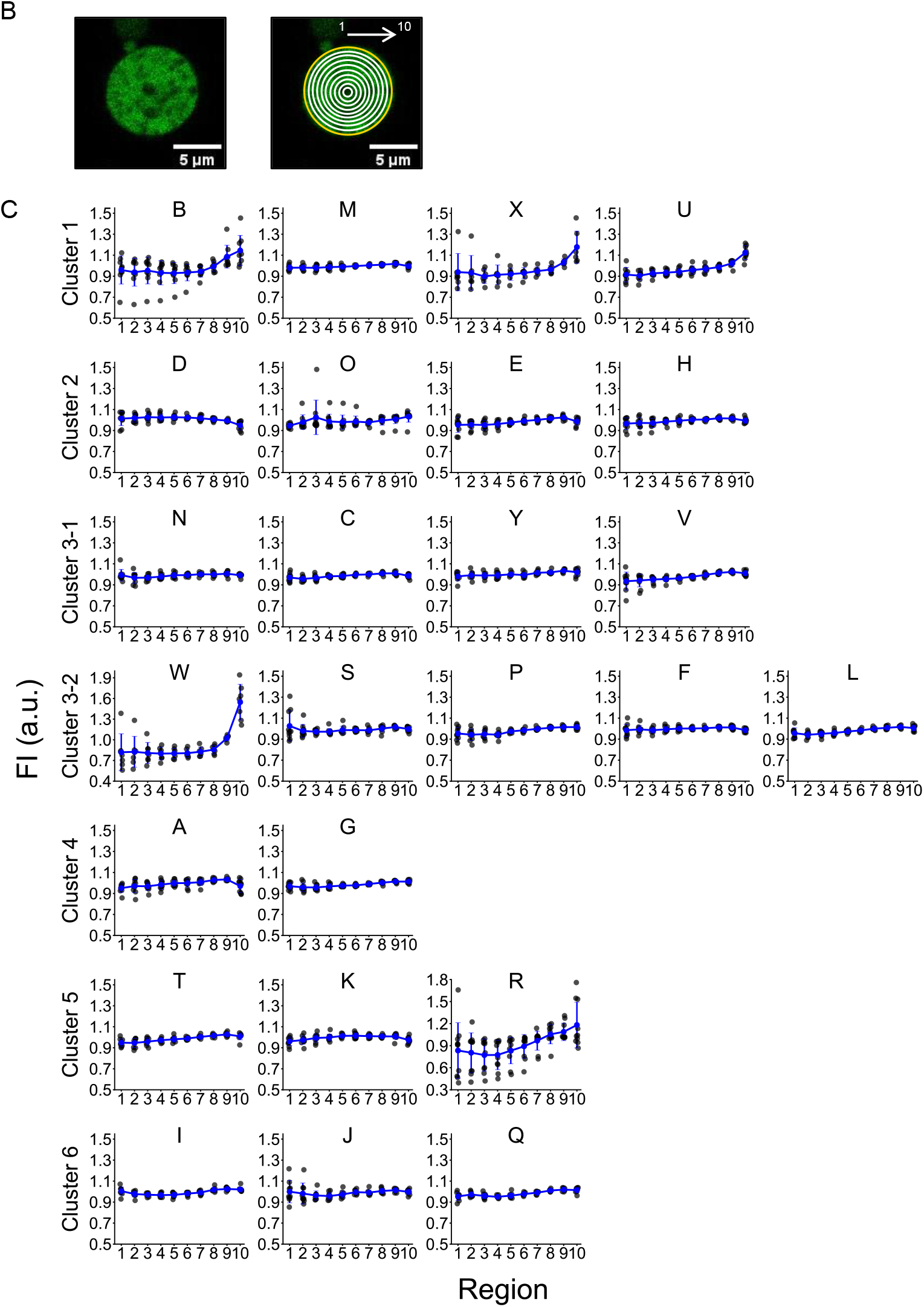
Subcellular localization of RasGEF-GFP. (A) Representative images of cells expressing RasGEF-GFP arranged by cluster (cluster 1 to cluster 6 from the top). Cluster 6 was excluded from the screening due to the weak expression of RBD_Raf1_-RFP. Scale bars, 10 μm. (B) 10 regions of the fluorescence intensity measurements. (C) Quantification of the subcellular localization of RasGEF. The blue circles and bars show the means and SDs, respectively, of more than 8 cells (black circles).

